# 2019 Association of Biomolecular Resource Facilities Multi-Laboratory Data-Independent Acquisition Study

**DOI:** 10.1101/2020.11.20.391300

**Authors:** Benjamin A. Neely, Paul M. Stemmer, Brian C. Searle, Laura E. Herring, LeRoy Martin, Mukul K. Midha, Brett S. Phinney, Baozhen Shan, Magnus Palmblad, Yan Wang, Pratik D. Jagtap, Joanna Kirkpatrick

## Abstract

Despite the advantages of fewer missing values by collecting fragment ion data on all analytes in the sample, as well as the potential for deeper coverage, the adoption of data-independent acquisition (DIA) in core facility settings has been slow. The Association of Biomolecular Resource Facilities conducted a large interlaboratory study to evaluate DIA performance in laboratories with various instrumentation. Participants were supplied with generic methods and a uniform set of test samples. The resulting 49 DIA datasets act as benchmarks and have utility in education and tool development. The sample set consisted of a tryptic HeLa digest spiked with high or low levels of four exogenous proteins. Data are available in MassIVE MSV000086479. Additionally, we demonstrate how the data can be analysed by focusing on two datasets using different library approaches and show the utility of select summary statistics. These data can be used by DIA newcomers, software developers, or DIA experts evaluating performance with different platforms, acquisition settings and skill levels.

## Background & Summary

Data-independent acquisition (DIA) is an alternative strategy to data-dependent acquisition (DDA) of MS2 fragmentation data in mass spectrometry. In DDA the instrument selects and fragments MS1 ions based on signal intensity. In DIA the mass spectrometer fragments analytes in predefined *m*/*z* windows. The MS2 data contribute to analyte identification and provide relative quantification. Both DDA and DIA approaches rely on sophisticated algorithms, and the interpretation of data is computationally intensive ^1–4^. Benefits of DIA include increased depth of coverage and between-sample uniformity (by to avoiding the stochastic nature of DDA acquisition), allowing unprecedented depth^5^ and speed^6^ of analysis. Recently, advances in instrumentation and algorithms have resulted in wider adoption^7–9^. In keeping with the mission of the Association of Biomolecular Resource Facilities (ABRF), the Proteomics Research Group (PRG) developed a multi-laboratory study (Figure 1), providing novice and expert users with samples and generic methods to benchmark their laboratories and empower participants to perform DIA.

**Figure 1.**
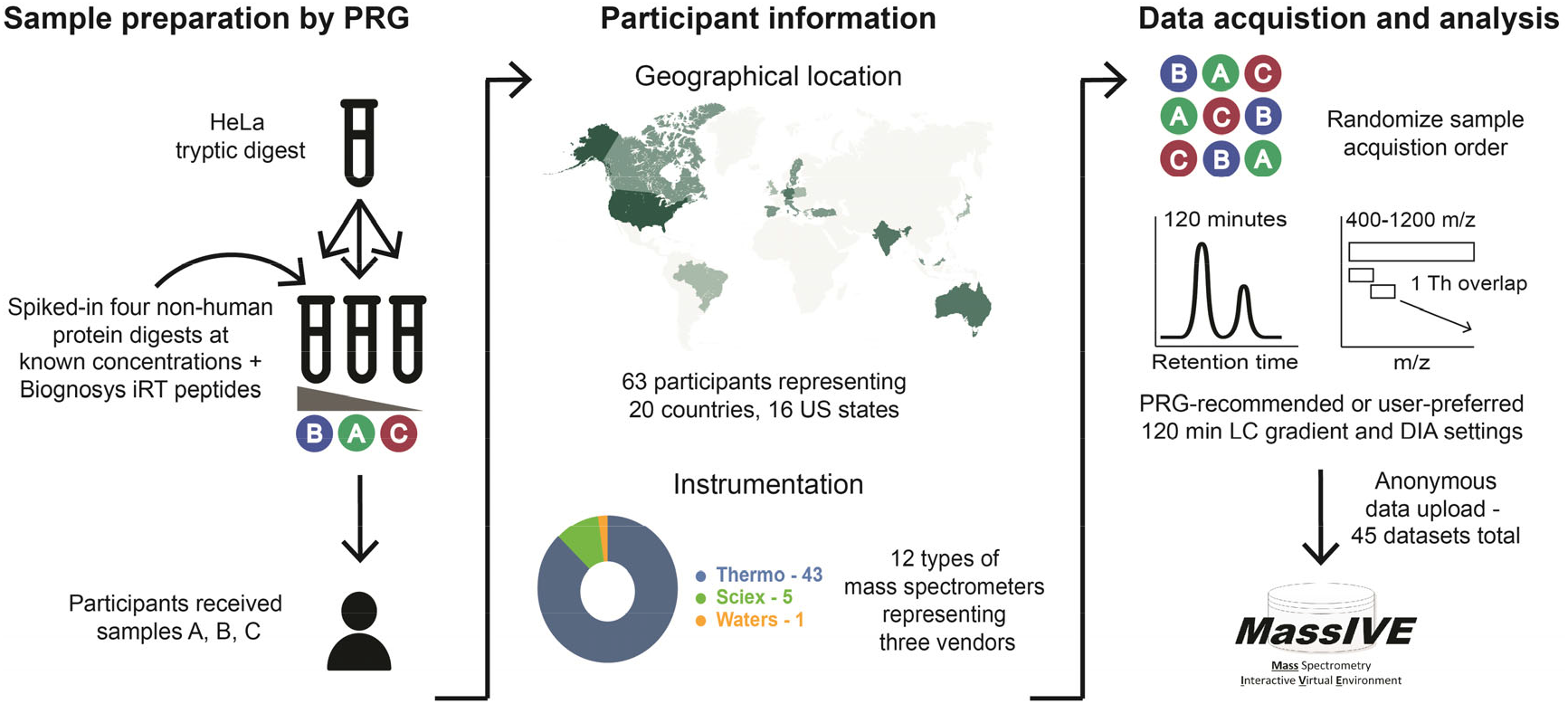
General description of study. The base sample in the study was a tryptic HeLa digest. Each participant received 20 ug of that sample as well as 20 ug of that sample with 2.5 or 10 fmol/μg of four non-human proteins: beta-galactosidase, lysozyme C, glucoamylase, and protein G. All samples also contained an internal retention time (iRT) peptide mix. Participants received dried samples identified only as A, B, or C. Participant identification was confidential and they could identify their dataset through a randomized identification number. Recommended LC and DIA settings were distributed with the samples though participants could use their preferred settings. Each lab was asked to run the samples in a specific randomized manner. Participants anonymously uploaded data to MassIVE, where it was curated by the PRG (renamed if needed and checked for file integrity) and re-uploaded to MassIVE MSV000086479.

Mixtures of proteomes^10,11^ have been used to benchmark proteomic workflows. We selected a set of four non-endogenous proteins in a human matrix^12^. The proteins: beta-galactosidase, lysozyme C, glucoamylase, and protein G were digested then spiked into the HeLa digest at 0, 2.5 and 10 fmol/μg (sample A - 2.5 fmol spike; sample B - 10 fmol spike, sample C was no spike). The four added proteins and two levels provided a wide range in signal intensities of the peptides, such that depth of spike-in coverage could reflect relative sensitivity between participants^13,14^. The study announcement was disseminated on the PRG website^15^, at conferences and via social media. Participants had varying prior experience and included mass spectrometers from different vendors (Figure 2; Table 1). A generic method (Supplemental File 1) was supplied and participants were asked to use a standard two-hour, two-step LC-gradient, a uniform static overlapping windowing strategy and a cycle time of approximately 3.5 s. Most participants followed these recommendations, (Table 2; Online-only Table 1). Of 63 laboratories that enrolled and received sample sets, 45 returned data. Some users had multiple instruments or compared different DIA methods, resulting in 49 datasets, 43 of which contain data for all replicates.

**Figure 2.**
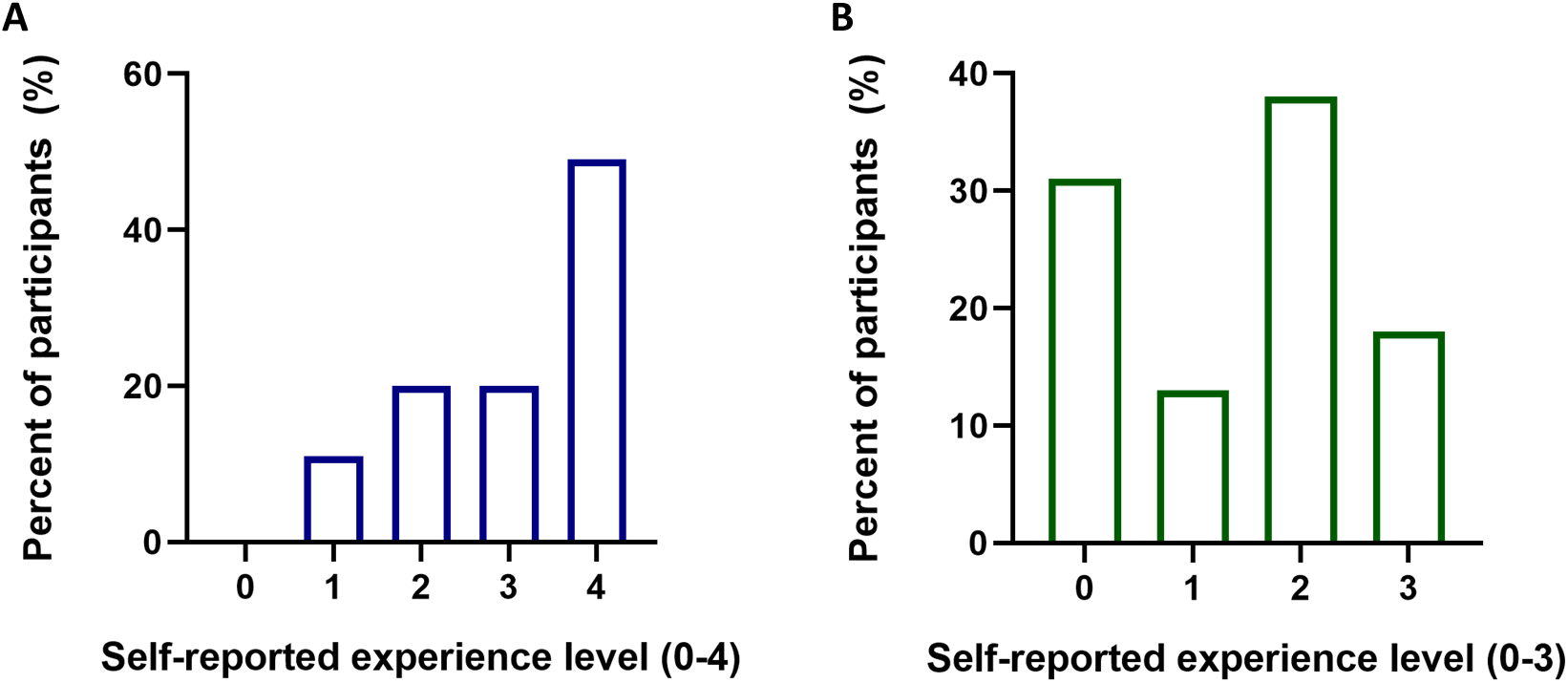
Self-reported experience level of 45 participants. An arbitrary scale was given to participants in a companion survey. **A. Self-reported experience LC-MS/MS level.** When asked about LC-MS/MS experience, participants were given the following choices: 0, “never set up myself”; 1, 0 to 2 years; 2, 3 to 5 years; 3, 6 to 9 years; 4, >10 years. **B. Self-reported DIA experience.** When asked about DIA experience participants were given the following choices: 0, Heard about it; 1, Tried it once; 2, Have done it a couple of times; 3, Expert.

**Table 1.**
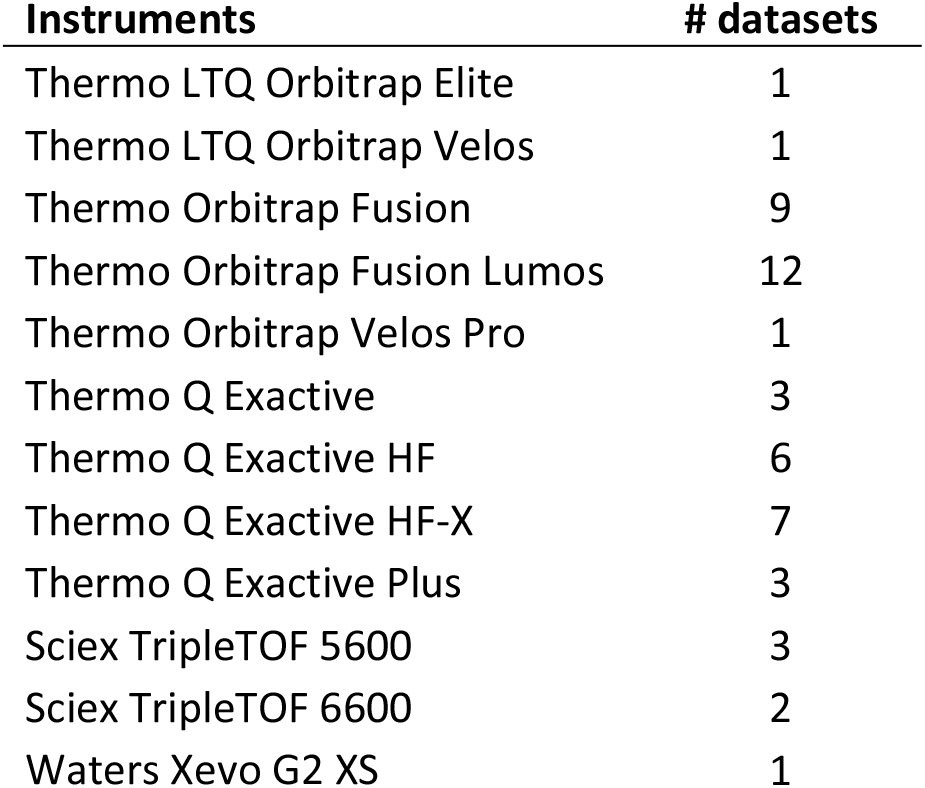
Instruments used in the study. The 45 participants deposited 49 datasets using 12 different instrument platforms from three different manufacturers.

| Instruments | # datasets |
| --- | --- |
| Thermo LTQ Orbitrap Elite | 1 |
| Thermo LTQ Orbitrap Velos | 1 |
| Thermo Orbitrap Fusion | 9 |
| Thermo Orbitrap Fusion Lumos | 12 |
| Thermo Orbitrap Velos Pro | 1 |
| Thermo Q Exactive | 3 |
| Thermo Q Exactive HF | 6 |
| Thermo Q Exactive HF-X | 7 |
| Thermo Q Exactive Plus | 3 |
| Sciex TripleTOF 5600 | 3 |
| Sciex TripleTOF 6600 | 2 |
| Waters Xevo G2 XS | 1 |

**Online-only Table 1.**
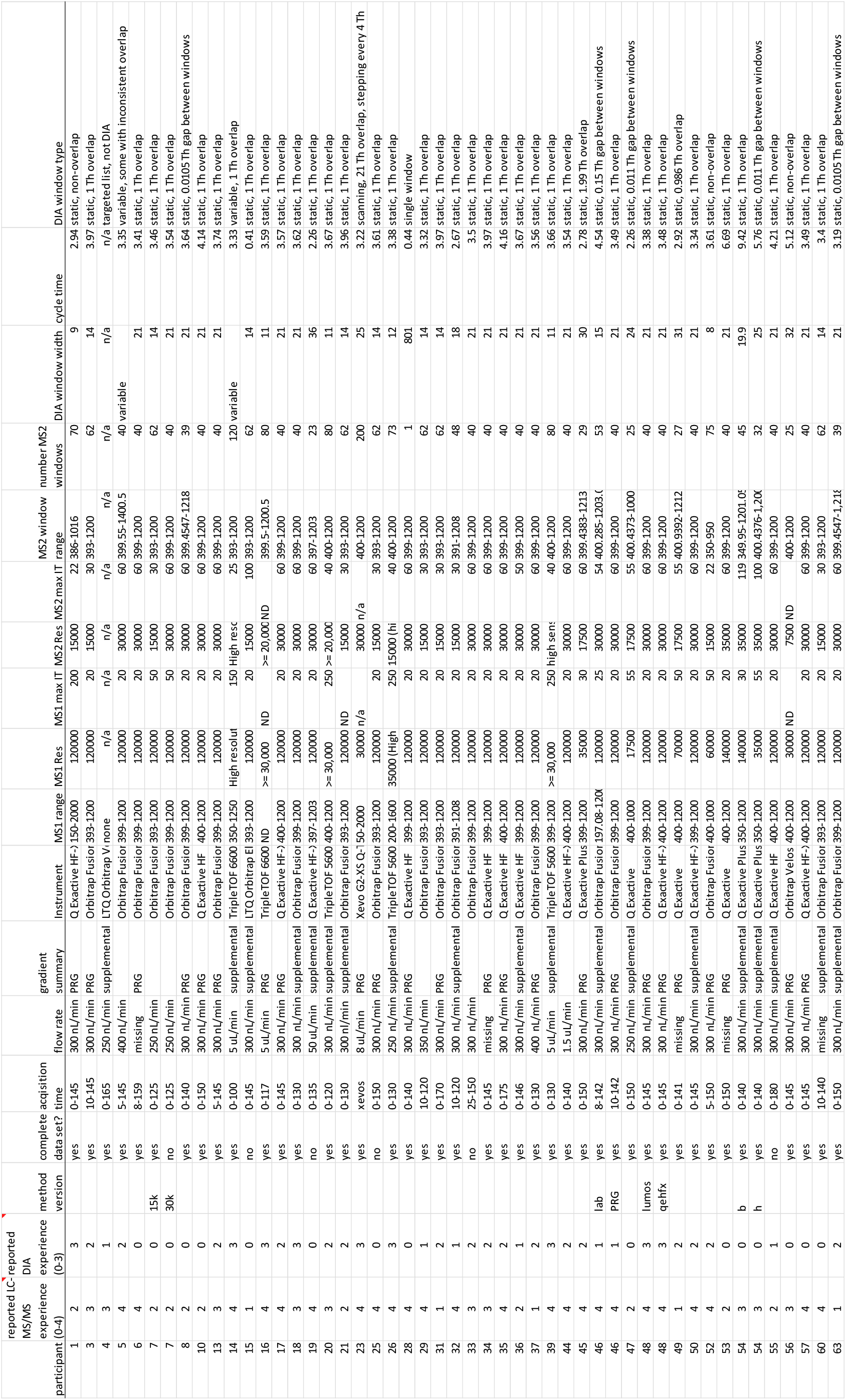
Meta information on 49 datasets. This table is available in MassIVE MSV000086479, but shown here for reference.

**Table 2.** Window scheme strategies used in the study. The DIA window strategies were divided into groups based on whether the DIA windows were static (i.e., the size did not change) and whether the DIA windows overlapped, while five datasets had gaps between the DIA windows.

| **window scheme** | **# datasets** |
| --- | --- |
| variable | 2 |
| static, non-overlap | 3 |
| static, w/overlap | 37 |
| static, w/gaps | 5 |
| single window | 1 |
| other* | 1 |

*\* used a targeted list*

**Table 3.**
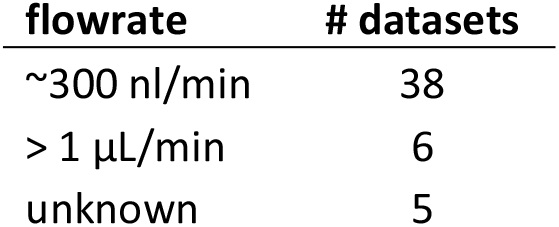
Flow rates used in the study. There were two main flow rates employed, either nL/min flow rates (approximately 300 nL/min; 250 to 400 nL/min), or μL/min flow rates (1.5 to 50 μL/min).

Other studies have used multi-laboratory DIA datasets to benchmark software tools^10,16^. Similarly, this new dataset has many uses, including benchmarking and user training. The range of user experience and instruments contributed to differences in data quality, providing a real-world dataset for evaluating how software normalization strategies are affected by data quality. Incorporation of known spikes facilitates evaluation of relative quantification using DIA ^17^. When the spiked proteins are ignored, each participant performed triplicate injections of experimental replicates, allowing for generation of useful summary statistics. Overall, this dataset provides opportunities for users to learn about acquisition methods and evaluate computational tools for DIA.

This dataset is also valuable to DIA software developers. The recommended acquisition method was not optimized for any platform, producing datasets conducted on different platforms with a similar acquisition strategy. All samples included iRT peptides and companion DDA and gas-phase fractionation data were also generated. Therefore, any software or library approach can use the data to evaluate and improve these approaches. The inclusion of the spike proteins in known amounts creates a unique opportunity to test new DIA strategies, such as MS1-based quantification^18^ and *in silico* generated libraries^19–23^.

In an initial analysis we have made a comparative analysis of data from two participants that used the same instrument. Each acquired additional data for library generation so it was possible to show how library construction and utilization effects results. Library strategy affected the number of proteins identified (Figure 3A-B), the precision of replicates (Figure 3A-B), and relative abundances of spike-in proteins (Figure 3C-D). Similar to reported observations^17^ these results highlight discrepancies when inferring protein abundance and the need to check relative quantification across the dynamic range. As these data and associated metadata are publicly available, we expect it to be used in benchmarking new tools and library strategies. Ongoing analysis of the dataset will provide more information and best-practice instrument settings for DIA, though the generic method provided performed unexpectedly well. With continued advancement of DIA methods, platforms optimized for DIA and improved computational strategies, this is an exciting time for the field, and we look forward to future multi-laboratory studies, enabling users and developers alike.

**Figure 3.**
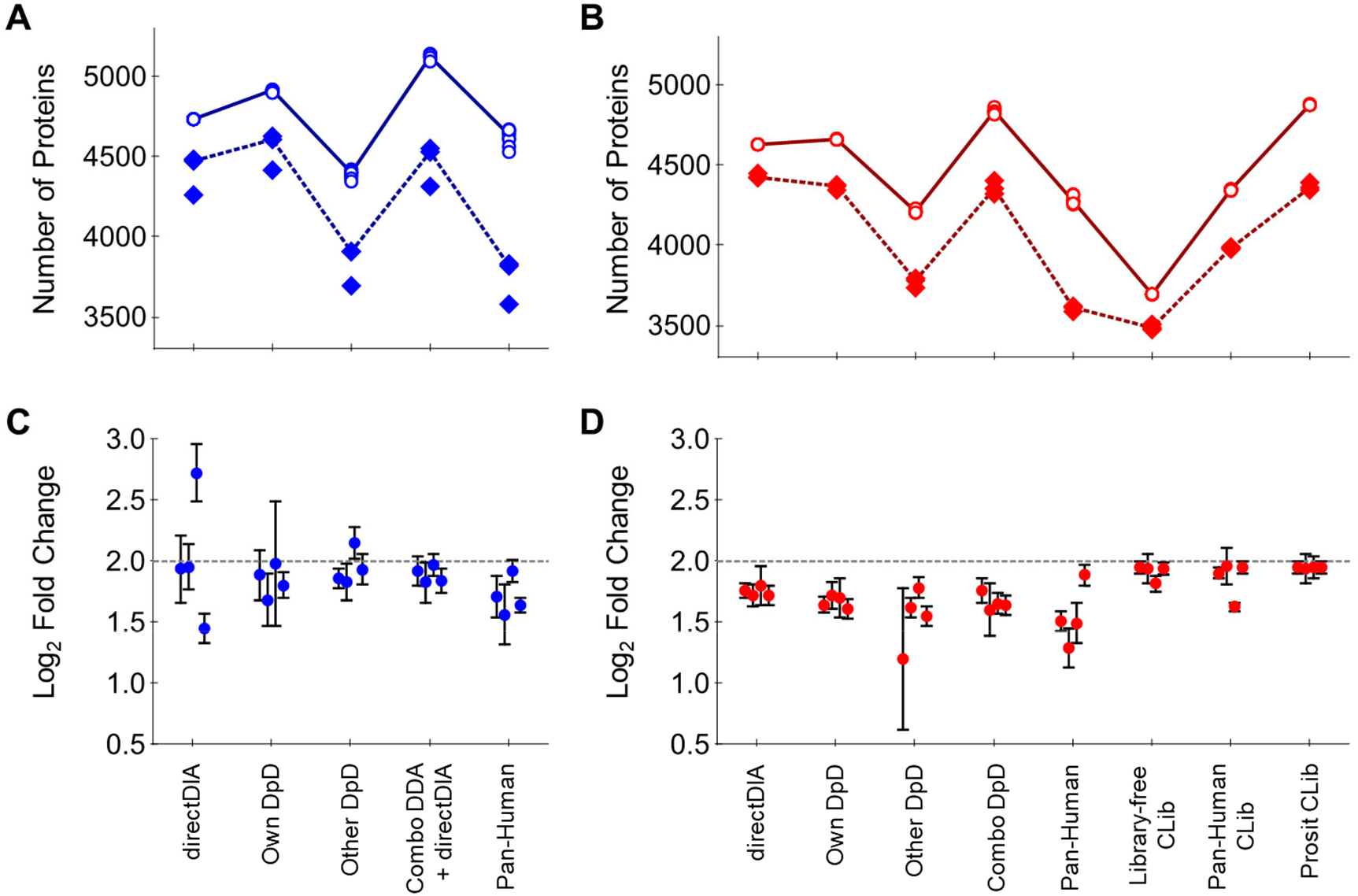
Comparison of library approaches with data from participants 3 and 48. Different library approaches were used to evaluate protein identification, technical variation and accurate relative quantification of four spike proteins with participant 48 **(A and C)** and participant 3 **(B and D)**. Eight library approaches are shown: directDIA, using the DIA files to generate the library; own **DpD** (DDA plus DIA), using the participant’s data to generate a library; other DpD, using the other participant’s DDA and DIA-based library; combined DpD, a combined library of the DDA and DIA data from both participants; Pan-Human, the Pan-Human library augmented with empirical evidence of the four non-endogenous spike-in proteins; Library-free CLib, chromatogram library; Pan-Human CLib, chromatogram library combined with Pan-human plus spikes; Prosit CLib, chromatogram library with prosit generated spectra. Only participant 3 generated a chromatogram library. **A-B.** Proteins identified using participant 48 data **(A)** or participant 3 data **(B)**. Hollow points are total identifications per sample with solid line being the average identifications. Solid points are number of proteins below 20 % CV with the dotted line being average proteins below 20 % CV. **C-D.** Estimated abundance of the four spike in proteins using the different library approaches for participant 48 **(C)** and 3 **(D)**. Log2 fold-change with 95% confidence intervals for the four spike-in proteins was determined between the 10 fmol/μg (sample B) and 2.5 fmol/μg (sample A) HeLa digest samples. The expected value was 2 (dotted line). Within each set of four points, left to right are ABRF-1, −2, −3 and −4, corresponding to the four spike-in proteins.

## Methods

### Study samples

Samples were prepared using Hela cells that were released from cell culture plates using trypsin. The cells were washed with PBS and cell pellets dispersed in MS-grade water then disrupted by sonication and diluted to a final protein concentration of 1 mg/mL. All digests were carried out using Promega trypsin with overnight incubations at 37 °C in 40 mM TEAB buffer and after reducing the sample with 10 mM DTT and alkylating with 30 mM IAA. Exogenous proteins were solubilized in MS-grade water and quantified from their absorbance spectra using calculated extinction coefficients ^24^. Equimolar amounts of the four proteins were combined prior to reduction and alkylation with DTT and IAA then digestion with trypsin. Digests of Hela and the exogenous protein mix were desalted using Oasis HLB (Waters) cartridges with a single step elution in 65 % (volume fraction) acetonitrile. The four exogenous proteins are: beta-D-galactosidase from *Escherichia coli* (Sigma, catalogue number G8511); Protein G from *Streptococcus aureus* (Sigma, catalogue number P4689); Lysozyme from *Gallus gallus* (Sigma, catalogue number L6876) and amyloglucosidase from *Aspergillus niger* (Sigma, catalogue number A7420). The digest of the exogenous proteins was added to the Hela lysate to achieve a concentration of 1 μM for each protein. This stock was diluted with the base HeLa digest to obtain 10, 2.5 or 0 fmol of the added proteins per 1 μg HeLa digest (sample A - 2.5 fmol spike; sample B - 10 fmol spike, sample C was no spike). Standard iRT peptides (Biognosys) were added to the HeLa plus exogenous protein mixtures. The three study samples of HeLa digest with exogenous proteins and iRT peptides were made one time and were aliquoted into 10 μg HeLa digests aliquots in 0.5 mL lo-bind tubes. These were dried by speed-vac and stored at −80 °C until shipped. Shipping was at ambient temperature.

### Study advertisement, enrolment, and timeline

Proteomics Research Group (PRG) members designed the study and announced it at the annual ABRF conference in April 2018. The study was also advertised at the annual conference of the American Society of Mass Spectrometry in June 2018. Interested participants contact details were collected via Google Survey and the distribution of samples began in September 2018 with the majority of participants receiving the samples by November 2018. Participating labs were located in 20 countries and 16 US States. The deadline for data return was extended to June 2019 to accommodate requests by some of the participants. Of the 63 participants who received samples, 45 labs returned datasets. Four participants performed multiple methods or acquired data on multiple instruments, resulting in 49 total datasets.

### Information given to participants

Each participant received a numerical study ID with the samples. The participants study ID was and is known only to that investigator and to the anonymizer. Documentation with information about study design, sample preparation, data acquisition and deposition was distributed electronically. This information is included in supplementary information (Supplemental File 1), though it has been edited from its original form to remove vendor contact information. The study documentation included suggestions for reconstituting the samples, LC gradient conditions and DIA data acquisition settings for the following platforms: Thermo Fusion and Fusion Lumos, Thermo QE-HFX, Sciex TripleTOF and Waters Xevo G2 XS platforms. Participants were encouraged to request guidance from members of the PRG if their platform was not included in the original guidelines. For those few investigators, a best attempt was made to design methods with approximately the same DIA cycle time. Finally, there were instructions on how to label the acquired data files and to complete and upload a survey that included self-reported metadata. Throughout the process participants were encouraged and given the means to remain anonymous even when securing technical assistance.

### PRG suggested sample resuspension and LC conditions

Participants received three dried samples that have been described. Participants using microflow received two complete sets of the three samples. The suggested method was to bring each up in 0.1 % formic acid, but did not specify the volume. It was expected that nanoflow systems would inject 1 to 2 μg on column, whereas microflow systems might require 4 to 8 μg on column. Participants had discretion to decide the appropriate injection amount for their system and to prepare the samples to allow for replicate injections.

Due to the diversity in LC systems and the latitude for participants to use either nano- or micro-flow applications, we relied on participants to design appropriate gradients that fit within basic guidelines. The suggestion for the study was a two-stage linear gradient lasting 110 to 130 min that we designated as the PRG gradient. The following was suggested: equilibration (trap or direct load) followed by a step from 5 to 25 % acetonitrile over 100 min, then 25 to 40 % acetonitrile over 20 min, and finally 40 to 90 % over 10 min. The final plateau could be held for 5 min before returning to 5 % acetonitrile over 1 min followed by re-equilibration. Roughly half, 23 of the 49, of the datasets reportedly used the PRG gradient (23 of 49 datasets), while 19 of the 49 datasets included specific gradient information that was deposited along with raw files on MassIVE. In general, a multi-step two-hour separation was performed by all participants. Participants were blinded to the sample identity so a run order of A, B, C, blank, B, C, A, blank, C, B, A was suggested to minimize systemic bias due to carryover.

### PRG suggested DIA conditions

When constructing a DIA experiment the MS2 mass range, number of MS2 windows, MS2 window width and time spent acquiring data all contribute to establishing the instrument’s cycle time, which is the time taken to scan all DIA windows one time, and therefore, how many data points are acquired during a peptide’s elution peak. For this study, we recommended static MS2 window widths covering 400 to 1200 *m/z*, with a 1 *m/z* overlap. As an example, this means that one window would stop at 420 *m/z* and the next window would start at 419 *m/z*. The majority of participants followed this recommendation (37 of 49 participants), but other strategies were selected by some participants (Table 2). The design of the study would produce a method with a 3.5 s cycle time. We assumed a 30 s peak width at base and, therefore, a 3.5 s cycle would produce between 7 and 10 data points per peak. We were aware that for participants with tighter peaks this would under sample. Overall, the parameters selected by participants largely achieved 3.5 s cycle (Online-only Table 1) as confirmed by evaluation of the window strategies using Skyline 4.2.0.19107 for each SA_R1 raw file for each submission. These are also reported in the windows.txt and windows.png files on MassIVE MSV000086479.

Specifying the instrument data acquisition time was difficult due to the diversity of platforms and instrument types. For example, for trap-based instruments the acquisition time is related to the transient time, AGC target, and max injection time. We provided general recommendations for QE-HFX, Fusion and Fusion Lumos and personalized recommendations for others, where given resolutions, with known transient times, and max injection times could be suggested to achieve a 3.5 s cycle. For non-trap based instruments such as the triple TOF line, it was much easier to specify instrument time since this is part of the method. In general, though, we suggested the following. For the Fusion and Fusion Lumos, 40 windows 21 *m/z* wide at 30 000 resolution or 62 DIA windows 14 *m/z* wide at 15 000 resolution. For the QE-HFX, 40 windows 21 *m/z* wide at 30 000 resolution. For tripleTOFs, 80 DIA windows 11 m/z wide. For specific settings such as max injection time and AGC (for trap based) or collision energy, please see Supplemental File 1.

#### Participant Actions

The actions and results of participants 3 and 48 will be discussed in detail.

### Participant 3 LC and DIA conditions

Participant 3 self-reported 6 to 9 years of LC-MS/MS experience, had performed DIA a couple times, and used a Thermo Fusion Lumos. The three samples were brought up in 20 μL 0.1% (volume fraction) formic acid to approximately 0.5 μg/μL. Peptide mixtures (2 μL injection; approximately 1 μg) were run in the order specified: A, B, C, blank, B, C, A, blank, C, B, A. The analysis was performed using an UltiMate 3000 Nano LC coupled to a Fusion Lumos mass spectrometer (Thermo Fisher Scientific) with a nano-ESI source. A trap/elute setup was used by trapping with a PepMap 100 C18 trap column (75 μm id x 2 cm length; Thermo Fisher Scientific) at 3 μL/min for 10 min with 2 % acetonitrile (volume fraction) and 0.05 % trifluoroacetic acid (volume fraction) followed by separation on an Acclaim PepMap RSLC 2 μm C18 column (75μm id x 25 cm length; Thermo Fisher Scientific) at 40 °C. Peptides were separated along the suggested PRG LC gradient except that the suggested 90 % acetonitrile (volume fraction) was not possible with the mobile phase setup used. Specifically, a 130 min gradient of 5 % to 32 % mobile phase B [80 % acetonitrile (volume fraction), 0.08 % formic acid (volume fraction)] over 100 min followed by a ramp to 50 % mobile phase B over 20 min and lastly to 95 % mobile phase B over 10 min at a flow rate of 300 nL/min.

Instrument acquisition settings for DIA were exactly those suggested for the 62 windows 14 *m/z* width at 15 000 fragment ion scan resolution (Supplemental File 1). Specifically, a default charge of 4 was used, no internal mass calibration was used, the ion funnel RF was 30 %, and full scan resolution of 120 000 (determined at 200 *m/z*), with an ion target value of 1.0×10^6^, and max injection of 20 ms. Full scan data was acquired from 393 to 1200 *m/z* in profile mode. For DIA settings, quad isolation was set at 14 *m/z* and a list of 62 mass centers was used to accomplish the suggested DIA window scheme, starting at 400 *m/z* and ending at 1193 *m/z*. This resulted in 62 DIA windows of 14 *m/z* width with 1 *m/z* overlap on edge of each window (ex. one window would stop at 420 *m/z* and the next would begin at 419 *m/z*). Fragmentation was performed using higher-energy collisional dissociation (HCD) at a normalized collision energy of 32. Fragmentation profile data was collected from 200 to 2000 *m/z* at 15 000 resolution. The max injection time was 30 ms and an ion target value 1.0×10^6^, and inject parallelizable ions was set to off. Data were acquired under Tune version 2.1 in XCalibur 4.0.

### Participant 48 LC and DIA conditions

Participant 48 self-reported >10 years LC-MS/MS experience, expert in DIA, and used a Thermo Fusion Lumos. The three samples were brought up in 20 μL 0.1 % (volume fraction) formic acid to approximately 0.5 μg/μL. Peptide mixtures (approximately 1 μg) were run in the order specified: A, B, C, blank, B, C, A, blank, C, B, A. The analysis was performed using an Nano Acquity (Waters) coupled to a Fusion Lumos mass spectrometer (Thermo Fisher Scientific). A trap/elute setup was used by trapping with a trap column (nanoAcquity Symmetry C18, 5μm, 180 μm × 20 mm) and an analytical column (nanoAcquity BEH C18, 1.7μm, 75μm × 250mm). The outlet of the analytical column was coupled directly to the MS using a Proxeon nanospray source. The peptides were introduced into the mass spectrometer via a PicoTip Emitter 360 μm OD × 20 μm ID; 10 μm tip (New Objective) and a spray voltage of 2.2 kV was applied. The capillary temperature was set at 300 °C. Mobile phase A was water with 0.1 % formic acid (volume fraction) and mobile phase B was acetonitrile with 0.1 % formic acid (volume fraction). The samples were loaded with a constant flow of mobile phase A (5 μL/min) onto the trapping column. Trapping time was 6 minutes. Peptides were eluted via the analytical column with a constant flow of 300 nL/min with the analytical column held at 40 °C. Peptides were separated along the suggested PRG LC gradient (Supplemental File 1).

Instrument acquisition settings for DIA were exactly those suggested for the 40 windows 21 m/z width at 30 000 fragment ion scan resolution (Supplemental File 1). Specifically, a default charge of 4 was used, internal mass calibration was used, the ion funnel RF was 30 %, and full scan resolution of 120 000 (determined at 200 *m/z*), with an ion target value of 1.0×10^6^, and max injection of 20 ms. Full scan data was acquired from 399 to 1200 m/z in profile mode. For DIA settings, quad isolation was set at 21 *m/z* and a list of 40 mass centers were used to accomplish the suggested DIA window scheme, starting at 409.5 *m/z* (center mass) and ending at 1189.5 *m/z* (center mass). This resulted in 40 DIA windows of 21 *m/z* width with 1 *m/z* overlap on the edge of each window. Fragmentation was performed using higher-energy collisional dissociation (HCD) at a normalized collision energy of 30. Profile data was collected from 200 to 2000 *m/z* at 30 000 resolution. The max injection time was 60 ms and an ion target value 1.0×10^6^, and inject parallelizable ions was set to True. Data were acquired under Tune version 2.1 in XCalibur 4.0.

### Participant 3 DDA and gas-phase fractionation runs

Participant 3 also performed additional analyses in order to provide data used for constructing spectral and chromatogram libraries. The remaining amounts (approximately 12 μL) of samples A (2.5 fmol spike) and B (10 fmol spike) were combined to obtain a solution that contained the spiked in proteins at approximately 6 fmol spike per μg HeLa digest. The same conditions were used as specified for DIA including the amount of sample injected and the gradient used. Data-acquisition settings were changed to standard DDA. For the data-dependent acquisition (DDA) runs, the Fusion Lumos was operated in positive polarity and data-dependent mode (topN, 3 s cycle time) with a dynamic exclusion of 60 s (with 10 ppm error). Full scan resolution using the orbitrap was set at 120 000 and the mass range was set to 375 to 1500 *m/z* collected in profile mode. Full scan ion target value was 4.0×10^5^ allowing a maximum injection time of 50 ms. Monoisotopic peak determination was used, specifying peptides and an intensity threshold of 1.0×10^4^ was used for precursor selection. Data-dependent fragmentation was performed using higher-energy collisional dissociation (HCD) at a normalized collision energy of 32 with quadrupole isolation at 0.7 *m/z* width. The fragment scan resolution using the orbitrap was set at 30 000, 110 *m/z* as the first mass, ion target value of 2.0×10^5^ and 60 ms maximum injection time and data type set to centroid.

To enable chromatogram library construction, “gas-phase fractionation” was performed^25^. The same injection volume and gradient were used. Five successive runs were performed using a staggered window approach described in detail by Searle *et al*.^25^ Briefly, a series of non-overlapping 4 *m/z* wide DIA windows are collected over a short enough mass range to maintain a reasonable DIA cycle. Then the cycle repeats but off-set by 2 *m/z*. This is repeated multiple times so that the full desired precursor mass range was covered. In the case of participant 3, there were five runs each with 2-cycles of 40 windows that were 4 *m/z* wide (detailed in Searle *et al.*^25^). The first run went 400 to 560 *m/z* then 398 to 558 *m/z*. The next four runs were 560 to 720 *m/z*, 720 to 880 *m/z*, 880 to 1040 *m/z* and 1040 to 1200 *m/z*. The raw file names were *TW1, *TW2, *TW3, *TW4, *TW5, respectively, shorthand for tight-window. For each run the instrument specifics were as follows: The Fusion Lumos was operated in positive polarity and no full scan data were acquired. Fragmentation was performed using higher-energy collisional dissociation (HCD) at a normalized collision energy of 32 with quadrupole isolation at 4 *m/z* width in conjunction with the two lists of 40 window centers. Fragment scan resolution using the orbitrap was set at 30 000 and the mass range was set to 200 to 2000 *m/z* collected in profile mode. The default charge was 4, RF lens was 30 % and the ion target value was 1.0×10^6^ allowing a maximum injection time of 60 ms.

### Participant 48 DDA runs (Lumos and QE)

Participant 48 also acquired additional data for library construction. The remaining parts of samples A (2.5 fmol spike) and B (10 fmol spike) were combined to obtain a solution that was approximately 6 fmol spike per μg HeLa digest. The same conditions were used as specified for DIA, including 1 μg injection and the same gradient, with the main change being data-acquisition settings. For the data-dependent acquisition runs, the Fusion Lumos was operated in positive polarity and data-dependent mode (topN, 3 s cycle time) with a dynamic exclusion of 15 s (with 10 ppm error). Full scan resolution using the orbitrap was set at 60 000 and the mass range was set to 375 to 1500 *m/z* collected in profile mode. Full scan ion target value was 2.0×10^5^ allowing a maximum injection time of 50 ms. Monoisotopic peak determination was used, specifying peptides and an intensity threshold of 5.0×10^4^ was used for precursor selection. Only multiply charged (2+ to 7+) precursor ions were selected for fragmentation. Isotopes were excluded. Data-dependent fragmentation was performed using higher-energy collisional dissociation (HCD) at a normalized collision energy of 30 with quadrupole isolation at 1.4 *m/z* width. The fragment scan resolution using the orbitrap was set at 15 000, 120 *m/z* as the first mass, ion target value of 2.0×10^5^ and 22 ms maximum injection time and data type set to centroid.

Participant 48 also performed additional analyses of the spike-in proteins alone to be included as a library when the data was searched against the Pan-Human library^26^. Participant 48 was supplied with a tryptic digest of approximately 16.9 pmol of each protein. The sample was resuspended in 170 μL (approx. 100 fmol/μL), the iRT kit added, and four injections were made in DDA and DIA, respectively, with the amount on column ranging from 100 fmol to 800 fmol (100, 200, 400, 800). These DDA and DIA runs of the spike-in proteins are described below.

The same conditions were used for the LC, with the exception that the nanoUPLC hardware was an M-Class NanoAcquity from Waters. The same gradient was applied (two-step PRG recommended), with the main change being the MS instrument settings for the QE-HFX (Thermo). For the data-dependent acquisition runs, the QE-HFX was operated in positive polarity and data-dependent mode (top15) with a dynamic exclusion of 20 s (with 10 ppm error). Full scan resolution using the orbitrap was set at 60 000 and the mass range was set to 350 to 1650 *m/z* collected in profile mode. Full scan ion target value was 3.0×10^6^ allowing a maximum injection time of 20 ms. Peptide setting was set to “preferred” and an intensity threshold of 1.0×10^4^ was used for precursor selection and AGC of 1.0×10^3^. Only multiply charged (2+ to 5+) precursor ions were selected for fragmentation. Isotopes were excluded. Data-dependent fragmentation was performed using higher-energy collisional dissociation (HCD) at a normalized collision energy of 27 with quadrupole isolation at 1.6 *m/z* width. The fragment scan resolution using the orbitrap was set at 15 000, 120 *m/z* as the first mass, ion target value of 2.0×10^5^ and 25 ms maximum injection time and data type set to profile. The default charge state was set to 2+. Data were acquired under Tune version 2.9 in XCalibur 4.0.

For the data independent acquisition runs, the same conditions as for the DDA were applied to the LC gradient. The following parameters were different for the QE-HFX acquisition: MS1 full scans were acquired using the orbitrap resolution set at 120 000 and the mass range was set to 400 to 1200 *m/z* and data were collected in profile mode. Full scan ion target value was 3.0×10^6^ allowing a maximum injection time of 20 ms. Data independent scans were set to 40 fixed windows, each of width 21 *m/z* (the same as for the Lumos DIA by participant 48). A maximum injection time of 60 ms was set with ion target value of 3.0×10^6^. Fragmentation (HCD) for the DIA scans in MS2 was carried out with a normalized collision energy of 30, the first MS2 mass was set to 200 *m/z* and data type set to profile. The default charge state was set to 3+.

### Analysis of participant 3 and 48 data

A preliminary analysis of the majority of participants data was presented at the ABRF 2019 meeting and is available for reference^27^. Herein we describe the analysis of two participants using Spectronaut (v13.6.190905.43655; Biognosys AG)^28^ and Scaffold DIA (v1.3.1; Proteome Software). Spectronaut and Scaffold DIA are two of the many available software packages capable of DIA analysis. They were selected for this project due to the expertise of the authors. We recommend similar analysis in other programs (see Data Usage), though the settings listed may not necessarily translate. Participants 3 and 48 both performed the replicate analysis of three samples using an Orbitrap Fusion Lumos. They both collected DDA in replicate of a combined sample, while data of just the digested spike proteins was only collected on a QE-HFX by participant 48. This allowed comparison of different library generation techniques within Spectronaut: directDIA, where only the DIA data is used; DpD, directDIA plus DDA where separate search archives (i.e., libraries) are constructed of the DIA data and the DDA, then combined; Pan-Human plus spikes, where a search archive of the spikes alone was combined with the Pan-Human library^26^. Since only Participant 3 collected data using gas-phase fractionation, this was used for generating a chromatogram library in Scaffold DIA and wasn’t utilized for Participant 48’s data.

The following settings were used in Spectronaut for directDIA libraries (setting tabs are bold), which can be retrieved as .xls and .kit files on MassIVE MSV000086479 as 03_lumos_directDIA and 48_lumos_directDIA. **Sequences:** Trypsin/P selected, max pep length 52, min pep length 7, two missed cleavages, KR as special amino acids in decoy generation, toggle N-terminal M set to true. **Labelling:** no labelling settings were used. **Applied modifications:** maximum of five variable modifications using fixed carbamidomethyl (C), and variable acetyl (protein N-term) and oxidation (M). **Identification:** per run machine learning, Q-value cut-off of 0.01 for precursors and proteins, single hits defined by stripped sequence, and do not exclude single hit proteins, PTM localization set to true with a probability cut-off of 0.75, kernel density p-value estimator. **Quantification**: interference correction was used with excluding all multi-channel interferences with minimum of 2 and 3 for MS1 and MS2, respectively. Proteotypicity filter was set to none, major protein grouping by protein group ID, minor peptide grouping by stripped sequence, major group quantity set to mean peptide quantity, a Major Group Top N was used (min of 1, max of 3), minor group quantity set to mean precursor quantity, a Minor Group Top N was used (min1, max of 3), quantity MS-level used MS2 area, data filtering by q-value, cross run normalization was used with global median normalization and automatic row selection, no modifications or amino acids were specified, best N fragments per peptide was set to between 3 and 6, with ion charge and type not used. **Workflow:** no workflow was used. **Post Analysis**: no calculated explained TIC or sample correlation matrix, differential abundance grouping using major group (from quantification settings) and smallest quantitative unit defined by precursor ion (from quantification settings), differential abundance was not used for conclusions but the following settings were used in the attached files: Student’s t-test, no group-wise testing correction, run clustering was set using the Manhattan distance metric and Ward’s method for linkage strategy and runs were ordered by clustering without Z-score transformation. The fasta files used are included in MassIVE MSV000086479, but briefly the UniProtKB Swiss-Prot 2018_06 human database (taxonomy:9606), canonical only was concatenated with the four spike proteins entries [ABRF-1 P00722 beta-galactosidase (*Escherichia coli*), ABRF-2 P00698 lysozyme C (*Gallus gallus*), ABRF-3 P69328 glucoamylase (*Aspergillus niger*), ABRF-4 Q54181 protein G’ (*Streptococcus* sp. group G)], the iRT Fusion sequence supplied by Biognosys [LGGNEQVTRYILAGVENSKGTFIIDPGGVIRGTFIIDPAAVIRGAGSSEPVTGLDAKTPVISGGPYEYRVEATFGVDESNAKTPVITGAPYEYRDGLDAASYYAPVRADVTPADFSEWSKLFLQF GAQGSPFLK], and a contaminants database of 247 entries. These two .fasta files are on MassIVE MSV000086479 as sp_human_180620_plus_PRG_ABRF_4_prot.fasta and contaminants_ 20120713.fasta, respectively.

For DpD approaches, the DDA data was used to generate a search archive with Pulsar using the same settings described for directDIA and the resulting search archive was combined with a library made directly from the DIA data using the settings described above. There were three DpD libraries created: each participant individually, then a combined library. These are included as 03_lumos_DpD, 48_lumos_DpD, and 03_lumos_48_lumos_DpD as .xls and .kit, on MassIVE MSV000086479. The Pan-Human search archive was downloaded within Spectronaut and is also available as Pan Human Library – ETH .xls and .kit on MassIVE MSV000086479. This was combined with a directDIA plus DDA library of digested spike proteins and is on MassIVE MSV000086479 as 48_qehfx_spikes_DpD .xls and .kit.

When searching with the DpD search archives or the Pan-Human derived archive, the following settings were used: **Data extraction:** maximum intensity extraction for MS1 and MS2, dynamic MS1 mass tolerance strategy with a correlation factor of 1, and a dynamic MS2 mass tolerance strategy with a correction of 1. **XIC extraction:** a dynamic XIC RT extraction window was used with a correction factor of 1. **Calibration:** allowed source specific iRT calibration with an automatic calibration mode, used a maximum intensity MZ extraction strategy, precision iRT was set to true with excluded deamidated peptides and a local (non-linear) iRT<->RT regression type, used Biognosys iRT kit, and no calibration carry-over. **Identification:** same settings as used in directDIA. **Quantification:** same as in directDIA. **Workflow:** no *in silico* library optimization, multi-channel workflow definition from library annotation with a fallback option as labelled, and no profiling or unify peptide peaks strategy was used. **Protein inference:** automatic. **Post analysis:** same settings as used in directDIA.

Since participant 3 also collected gas-phase fractionation data (described above), this DIA data was also processed in Scaffold DIA (v1.3.1) three different ways: (1) by creating a chromatogram library using only the gas-phase fractionation data, (2) using these data combined with the Pan-Human library and (3) using these data combined with a Prosit *in silico* library. The Pan-Human library was converted directly using EncyclopeDIA (v0.8.1)^25^ from phl004_canonical_sall_osw.csv, downloaded from the SwathAtlas repository (https://db.systemsbiology.net/sbeams/cgi/PeptideAtlas/GetDIALibs). The Prosit predictions used the EncyclopeDIA library generation defaults^20^. These defaults were predictions for +2H/+3H peptides between 396.4 and 1002.7 m/z with up to 1 missed cleavage. The NCE setting was 33, assuming all peptides were fragmented in DIA as +2H peptides. The additional four ABRF peptides were predicted using the same pipeline, but for all +2H/+3H/+4H/+5H peptides between 396.4 and 1002.7 *m/z* with up to 2 missed cleavage. Again, the NCE setting was 33, assuming all peptides were fragmented in DIA as +2H peptides. The resulting library files for Pan-Human and Prosit human plus PRG spike proteins can be found on MassIVE MSV000086479 as combined_prg_pan_human.dlib and combined_prg_sprothuman.dlib, respectively.

All raw data files were converted to mzML format (within Scaffold DIA) using ProteoWizard (v3.0.18342)^29^. In the first case where an external library was not used, the reference spectral library was created by EncyclopeDIA (v0.8.1)^25^. Reference samples were individually searched against the same fasta described above, with a peptide mass tolerance of 10.0 ppm and a fragment mass tolerance of 10.0 ppm. Fixed modifications considered were: Carbamidomethylation C. In the second case, when combining the data with the Pan-human library, the reference spectral library files were individually searched against a combined fasta and the dlib with a peptide mass tolerance of 10.0 ppm and a fragment mass tolerance of 10.0 ppm. Variable modifications considered were: Oxidation M and Carbamidomethylation C (since these are used in the Pan-Human library). In the third case, when combining the data with Prosit predictions, the reference spectral library files were individually searched against a combined fasta and the dlib with a peptide mass tolerance of 10.0 ppm and a fragment mass tolerance of 10.0 ppm. Variable modifications considered were: Carbamidomethylation C.

For all three search approaches the digestion enzyme was assumed to be trypsin with a maximum of 1 missed cleavage site allowed. Only peptides with charges in the range +2 to +3 and length in the range 6 to 30 were considered. Peptides identified in each search were filtered by Percolator (v3.01.nightly-13-655e4c7-dirty)^30–32^ to achieve a maximum FDR of 0.01. Individual search results were combined and peptides were again filtered to an FDR threshold of 0.01 for inclusion in the reference library.

Analytical samples (i.e., the replicate injections of the three ABRF samples participant 3 analyzed), were aligned based on retention times and individually searched against 03 Chromatogram Library.elib, 03 - PH plus CL library.elib or 03 - Prosit plus CL library.elib (created as described above and available on MassIVE MSV000086479) with search settings identical to those used to create the reference library. Peptide quantification was performed by EncyclopeDIA (v0.8.1)^25^. For each peptide, the five highest quality fragment ions were selected for quantitation. Proteins that contained similar peptides and could not be differentiated based on MS/MS analysis were grouped to satisfy the principles of parsimony. Proteins with a minimum of two identified peptides were thresholded to achieve a protein FDR threshold of 1.0 %. These files are available as 03 - CL only.sdia, 03 - PH plus CL.sdia and 03 - Prosit plus CL.sdia.

For all approaches, intensity values of the four spike-in proteins were used to compare relative quantification between the different approaches. Specifically, Sample A versus Sample B was used to evaluate how well each approach measured the predicted 4-fold difference in protein concentration. To easily calculate the 95% confidence interval of each fold-change the topTable function within the limma package (v3.40.6)^33^ in R (v3.6.0; 64-bit) was used with the argument “confint=TRUE”. To accomplish this, first exported intensity values were transformed to log2 values and this matrix was used with limma. A summary figure outlining the workflow for the data from participants 3 and 48 is shown in Figure 4.

**Figure 4.**
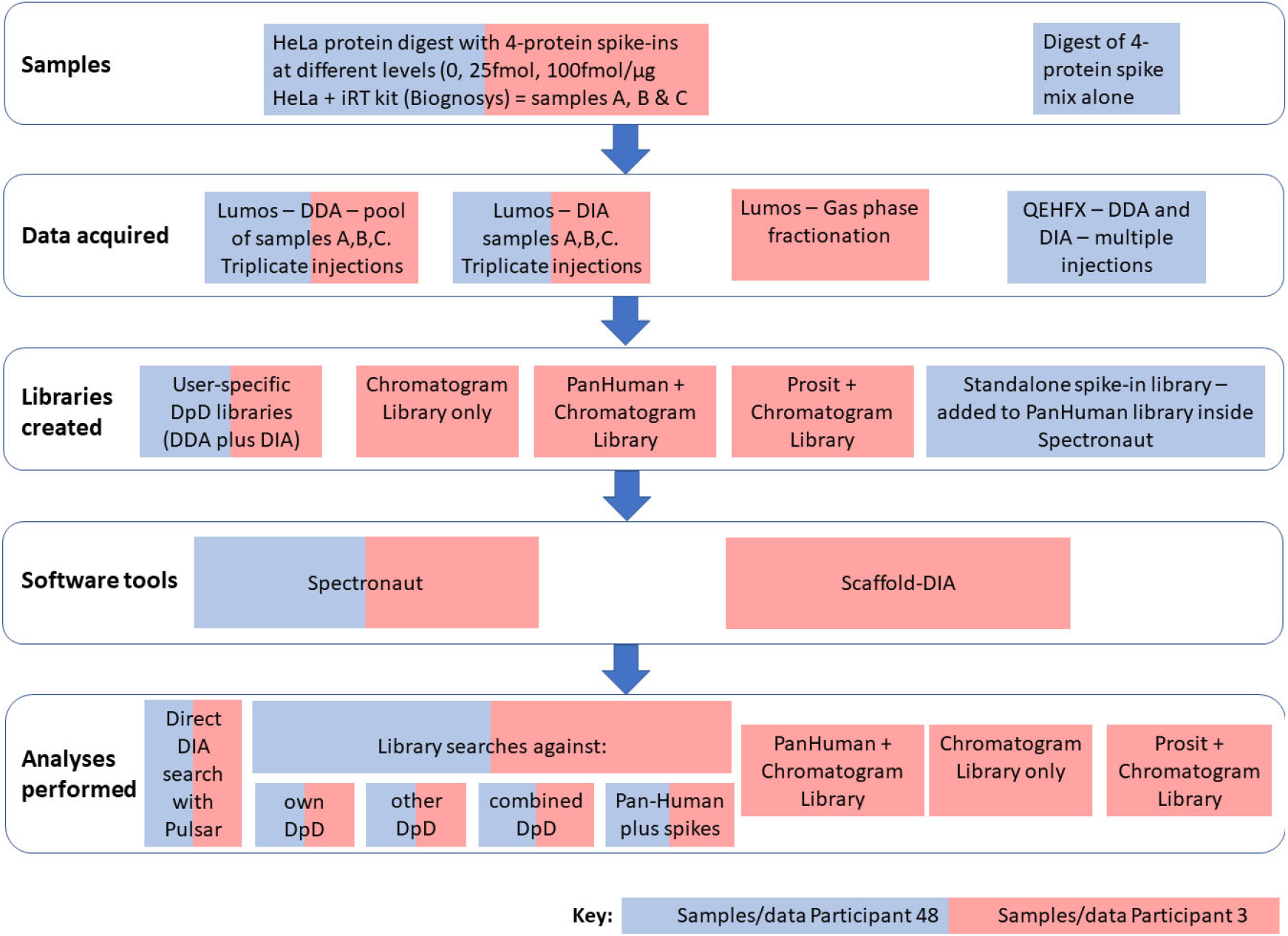
Summary flowchart showing steps taken from samples assigned to participants 3 and 48.

## Data Records

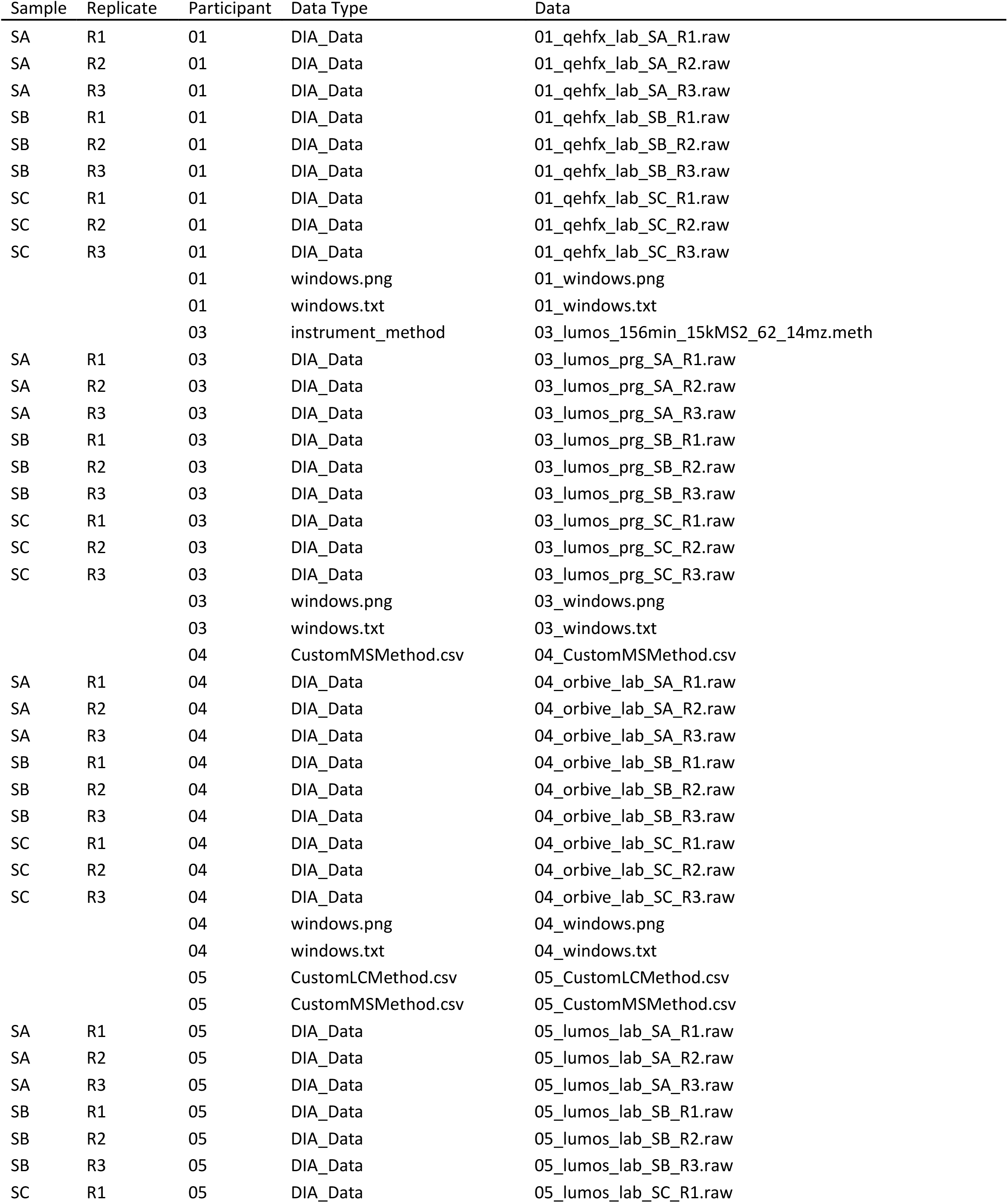
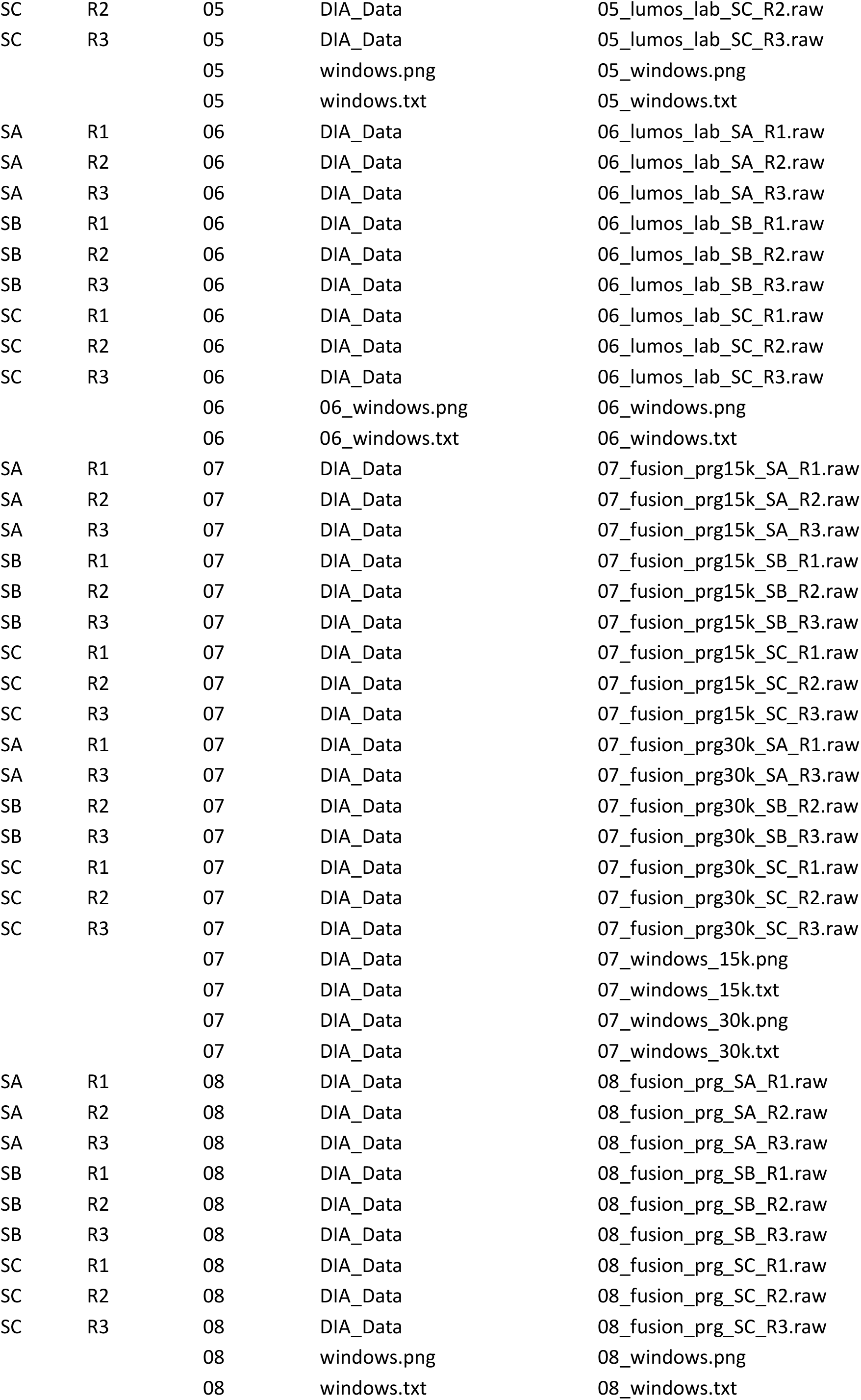
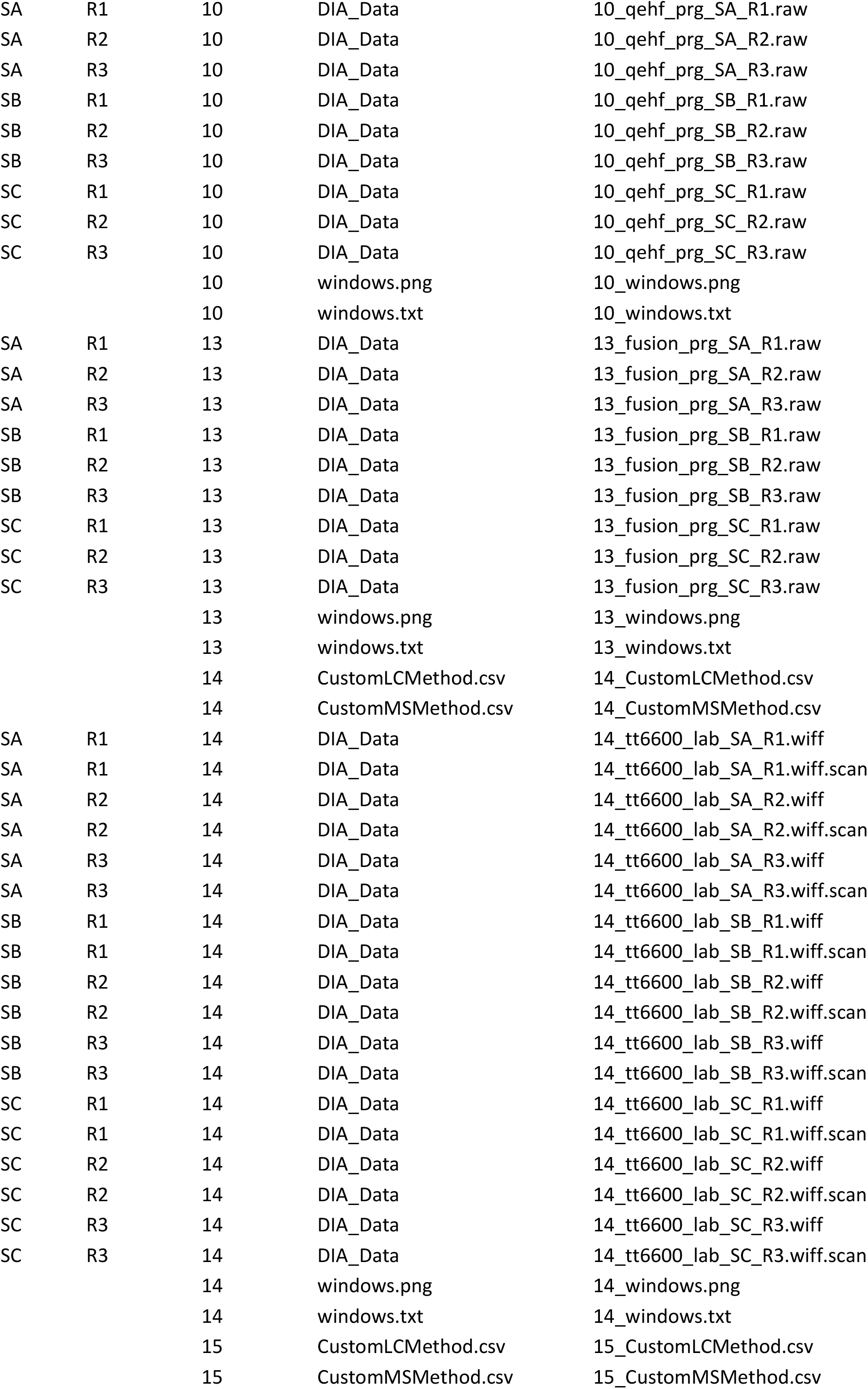
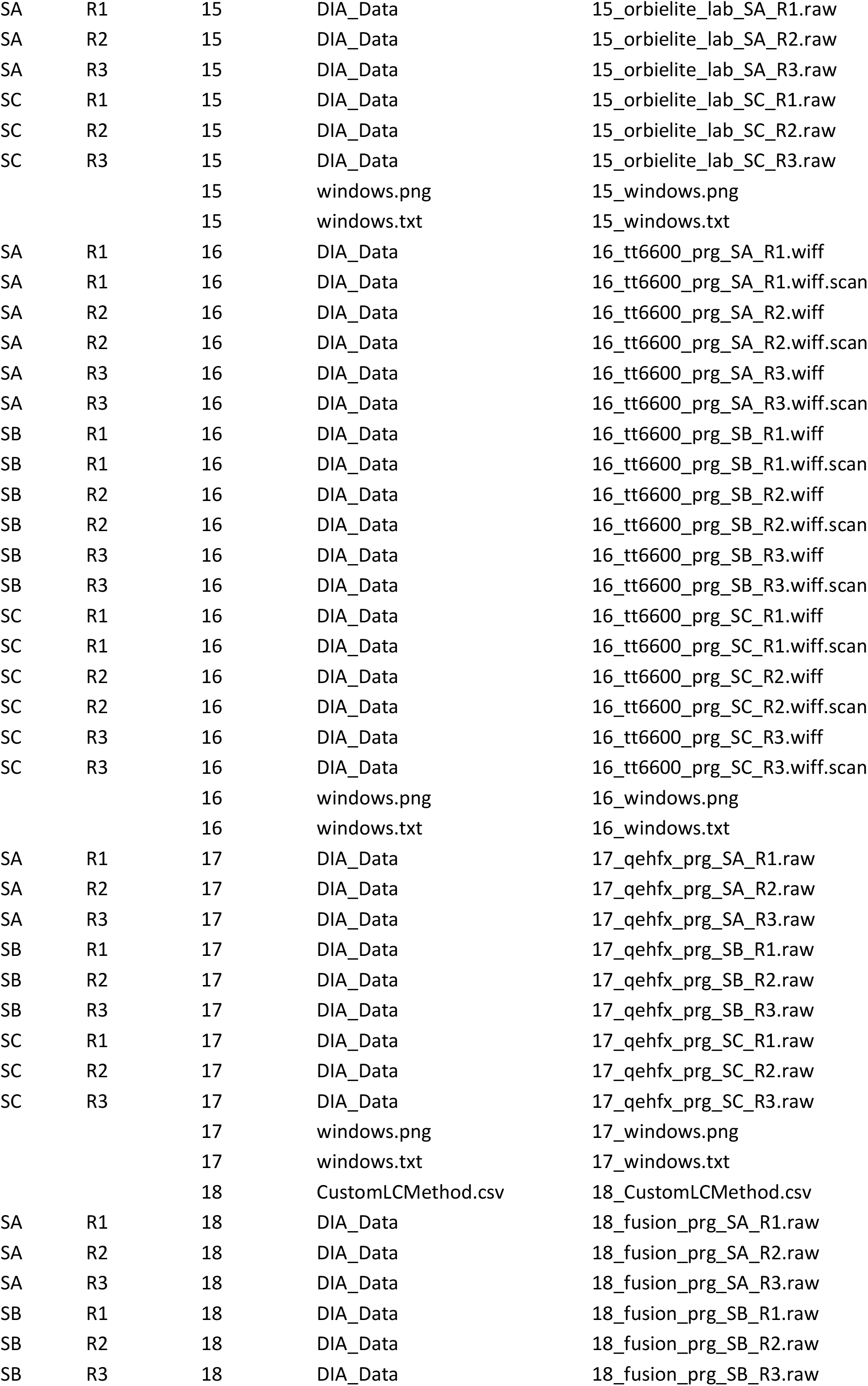
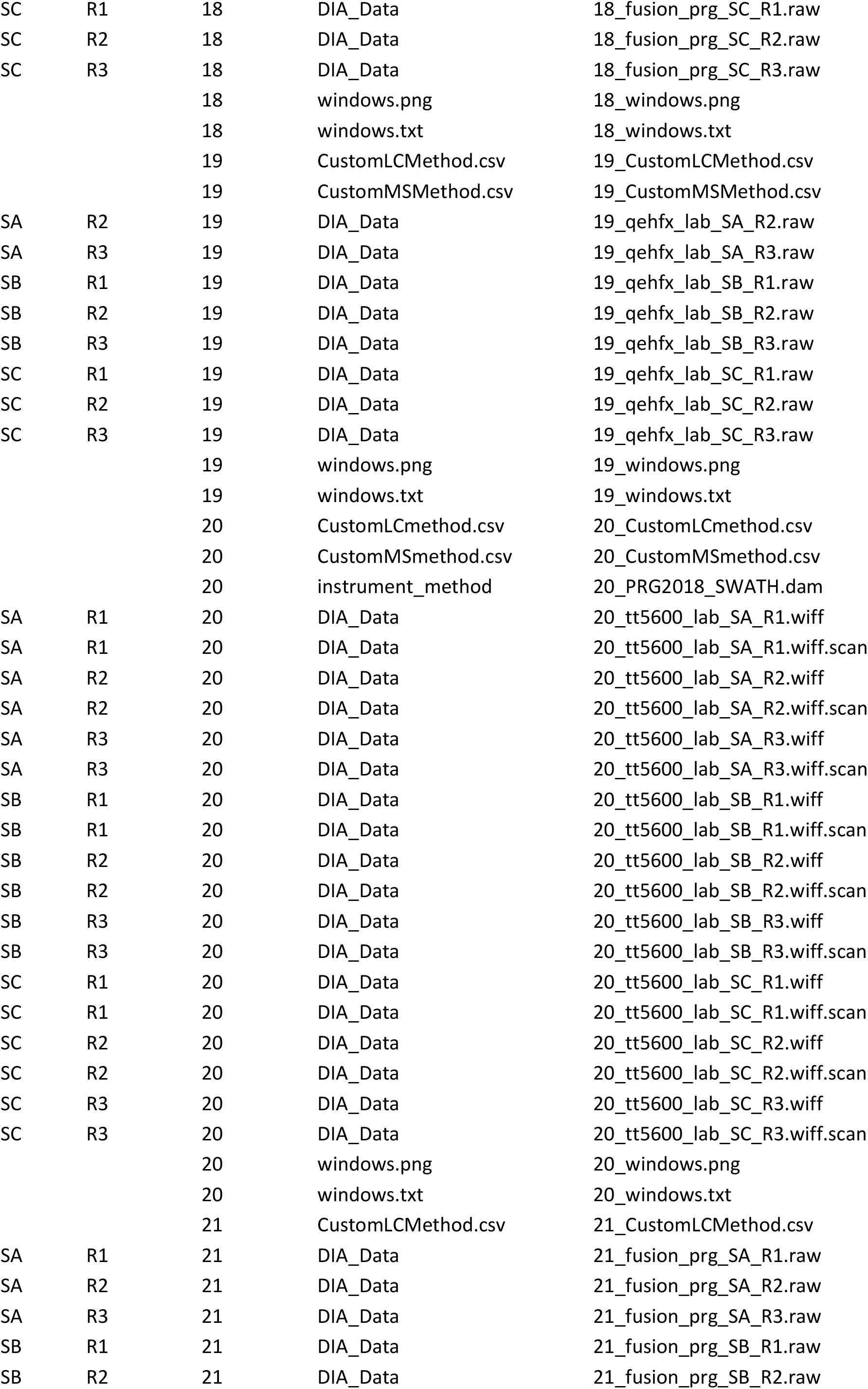
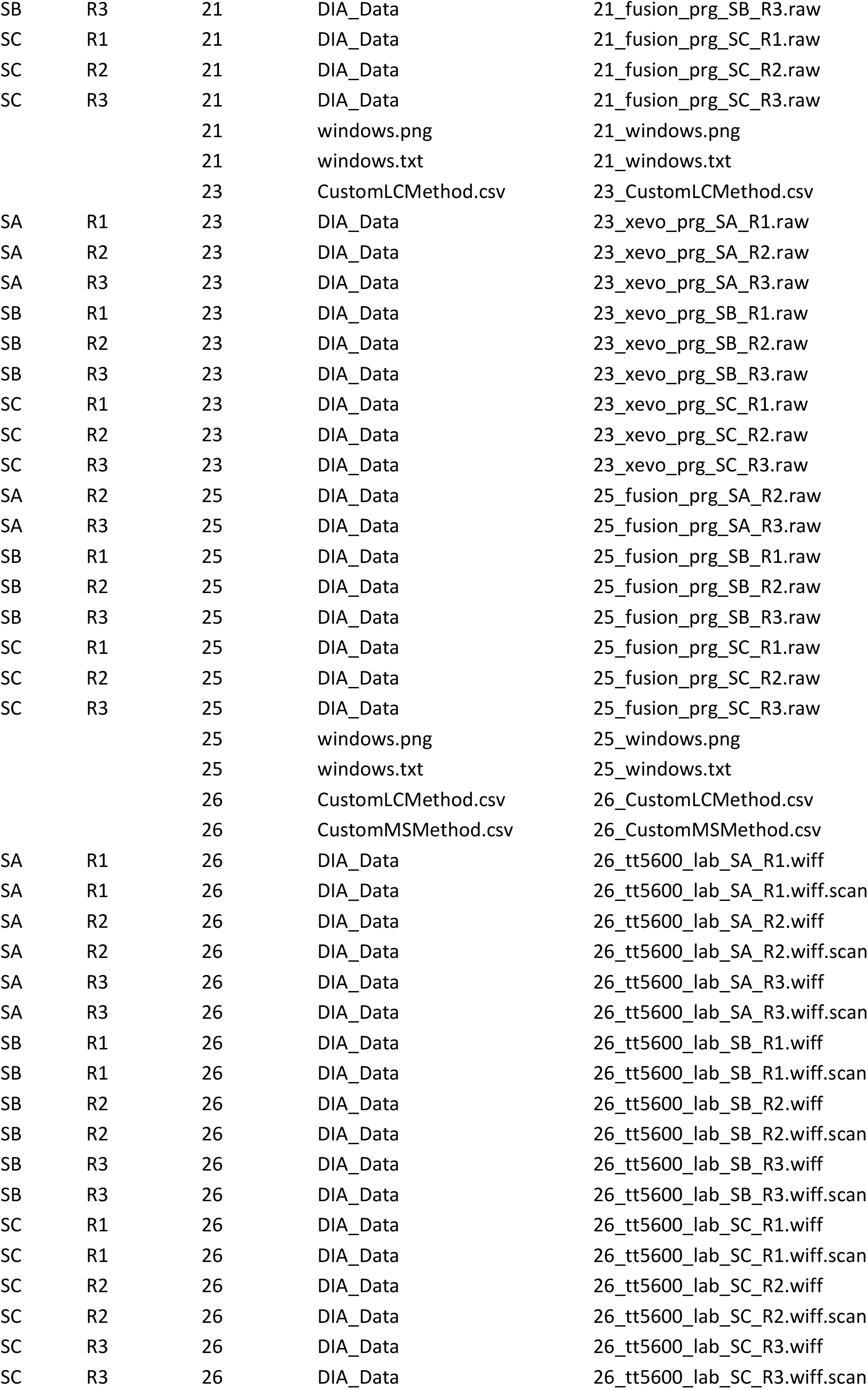
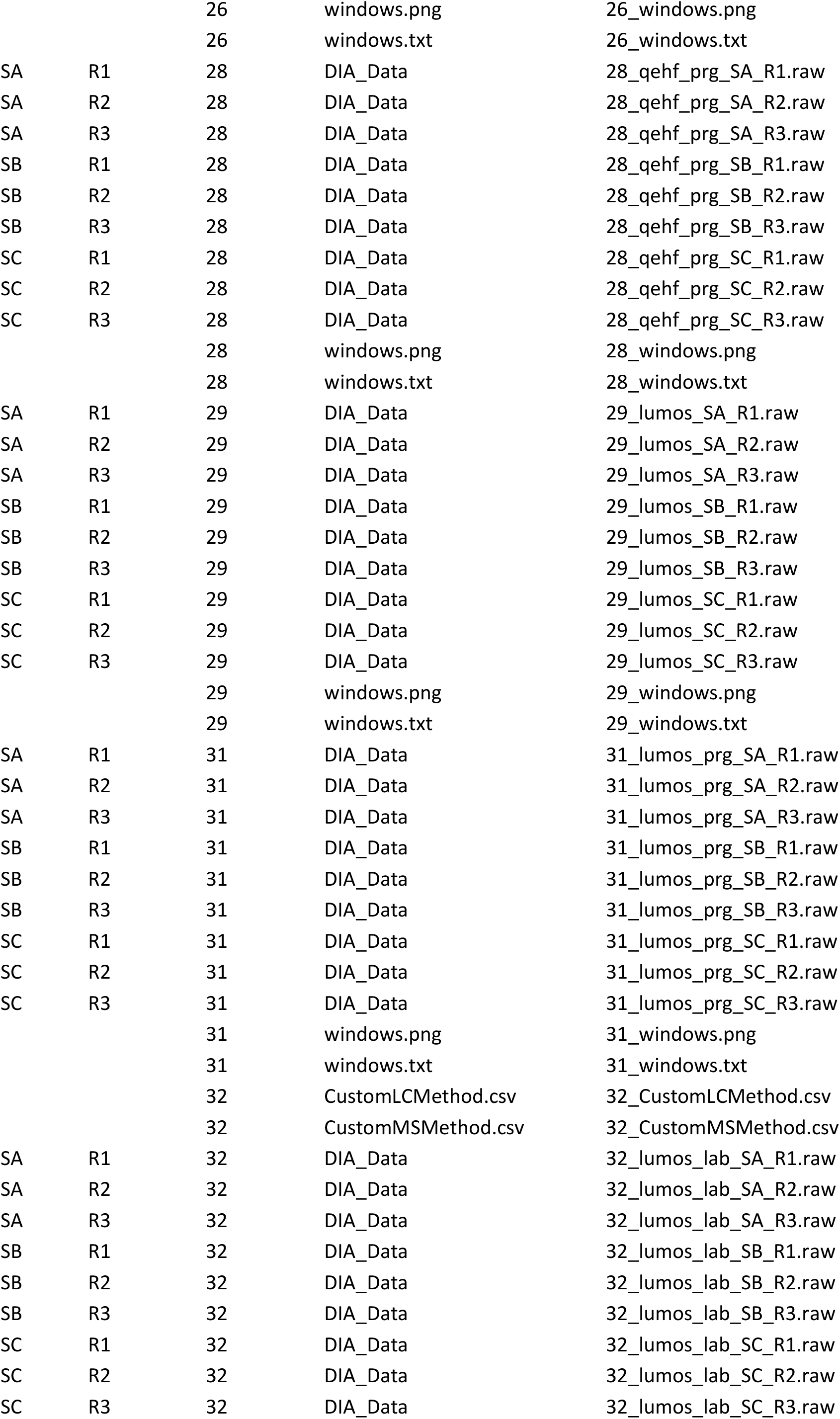
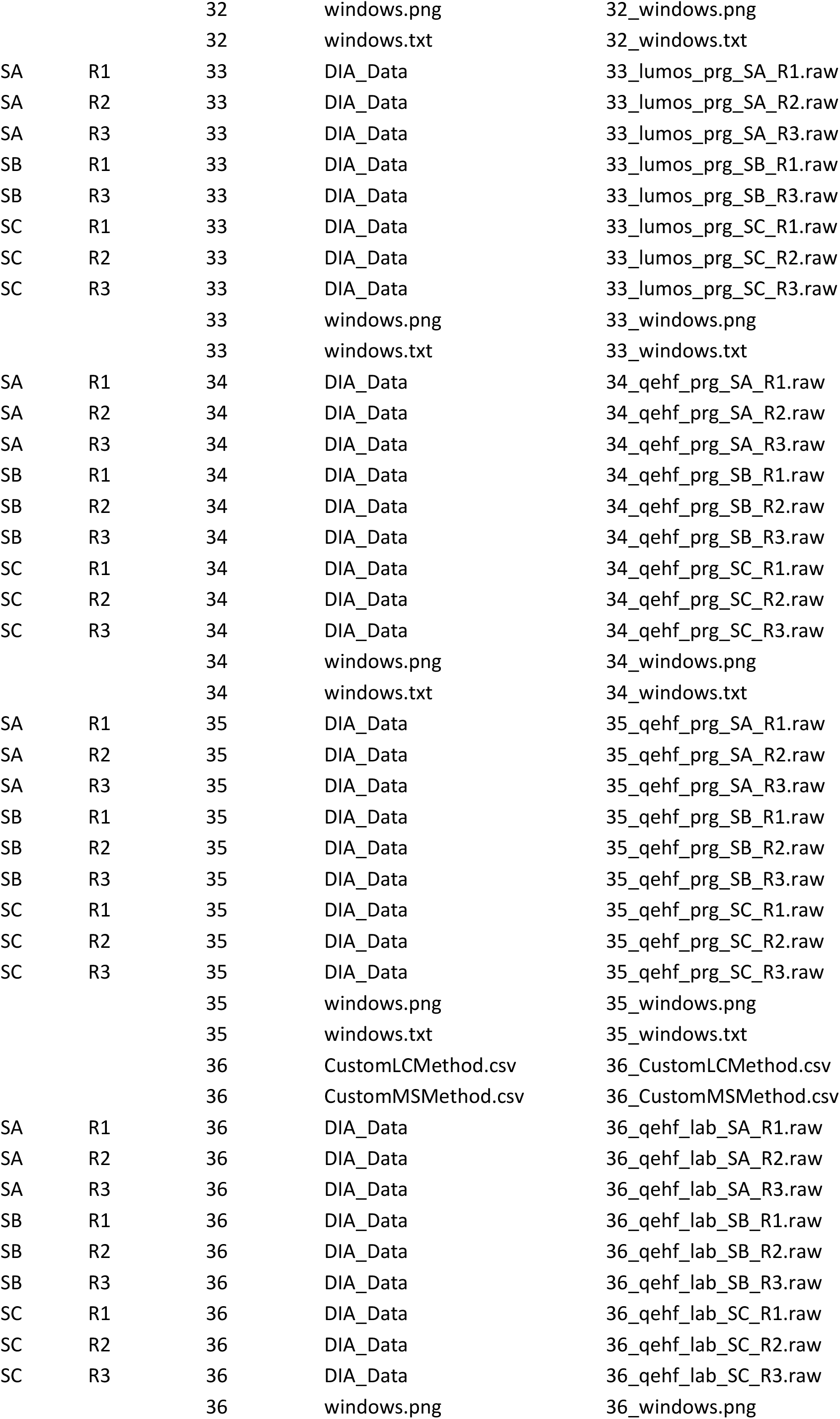
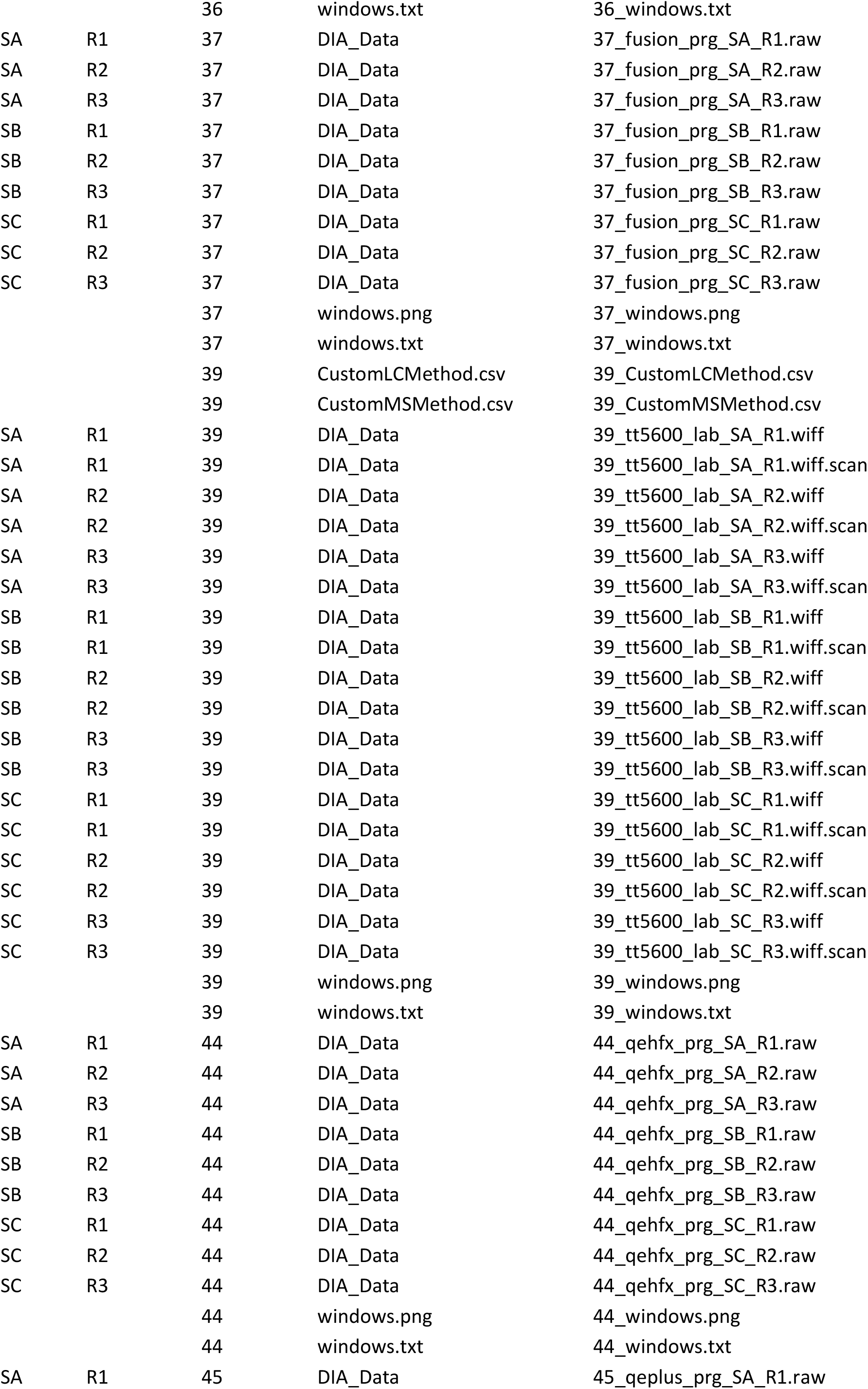
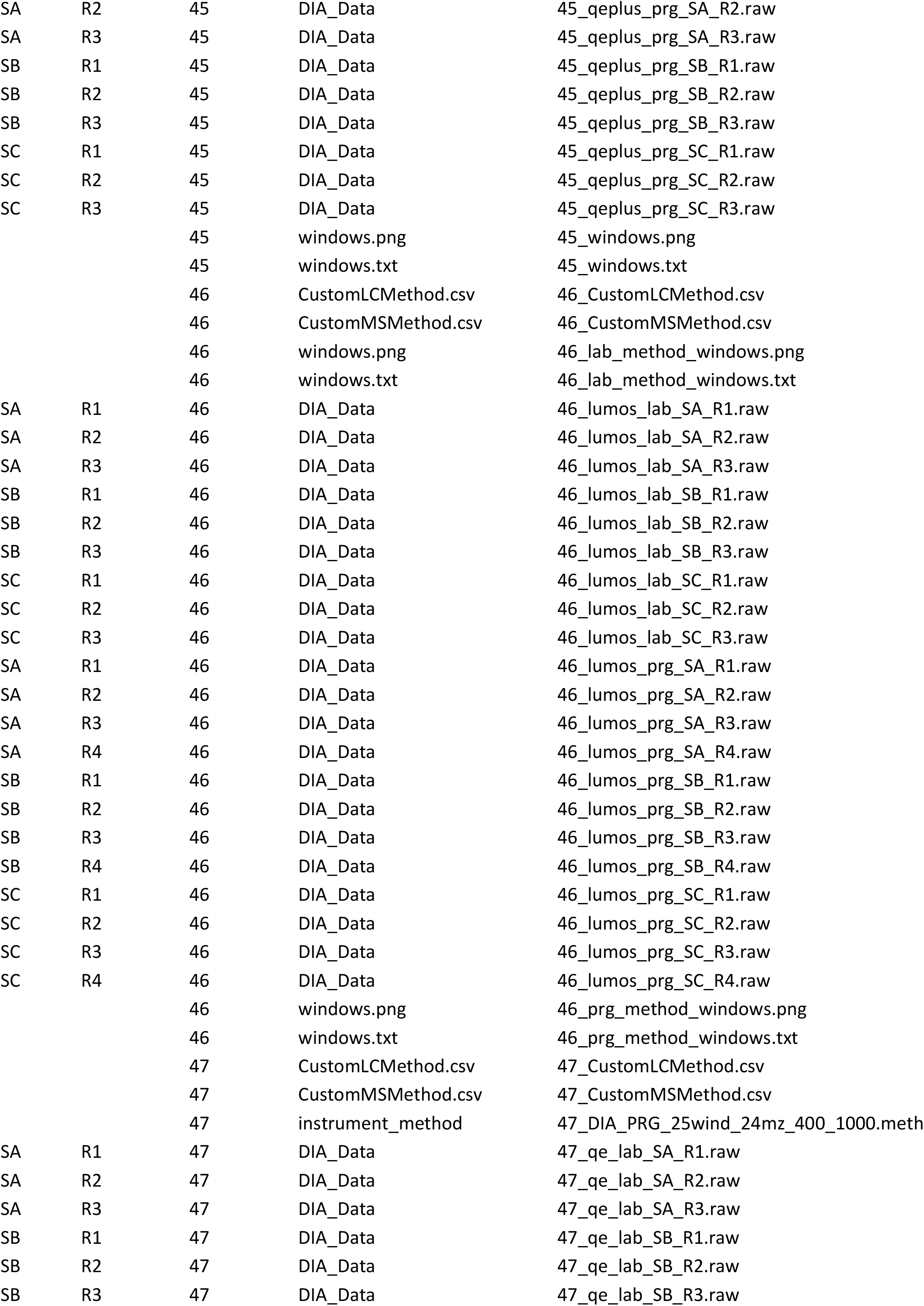
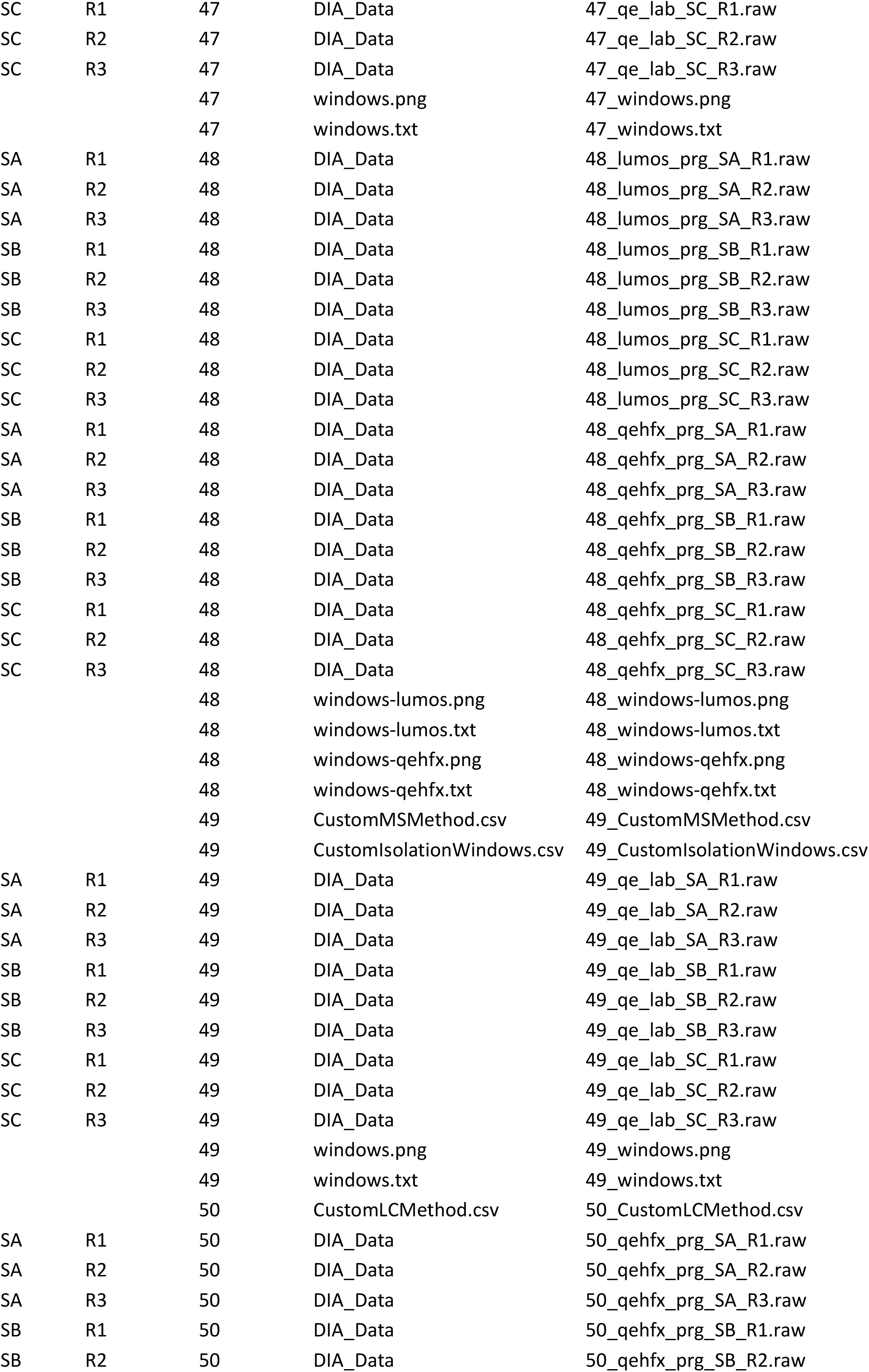
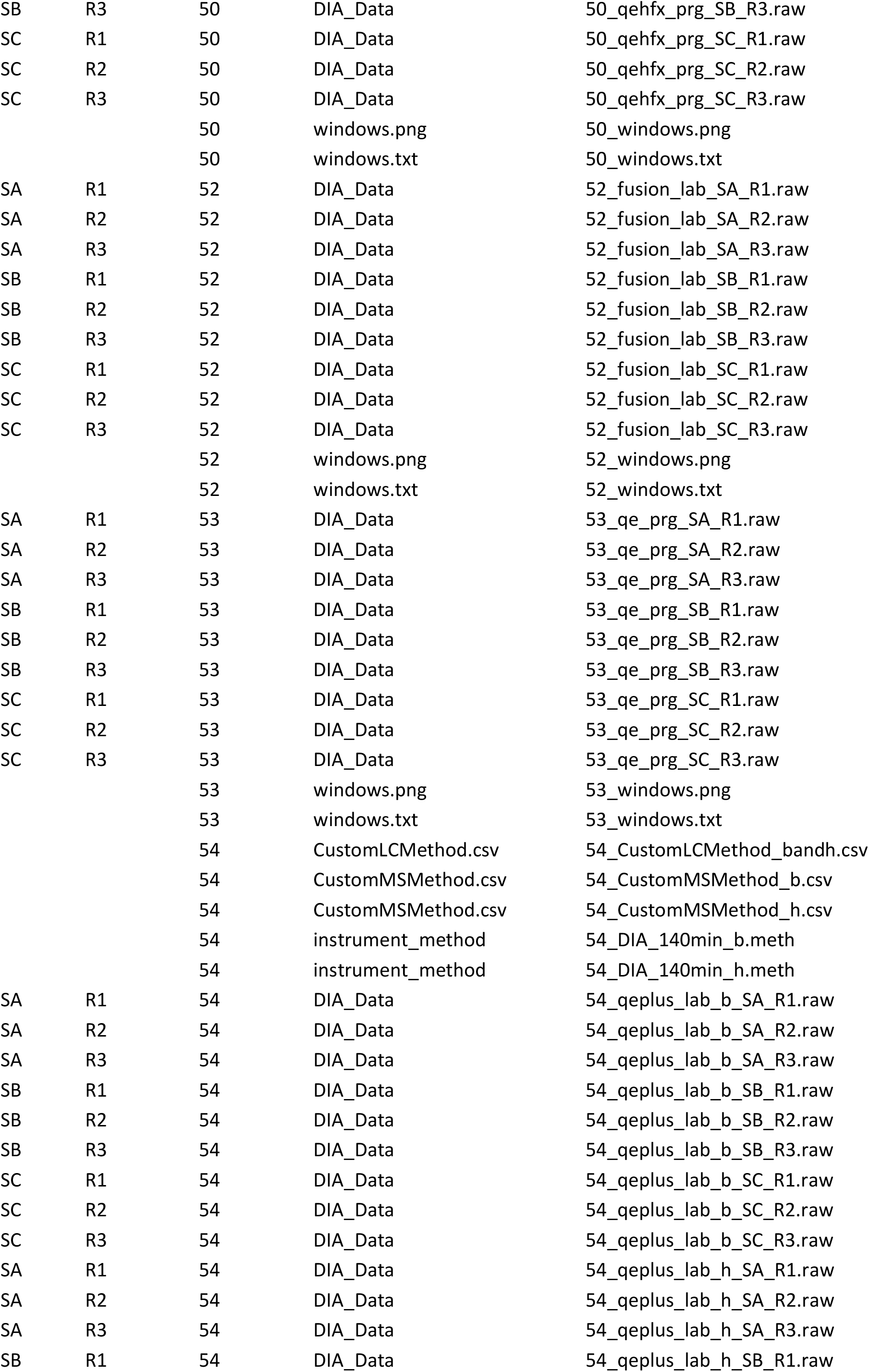
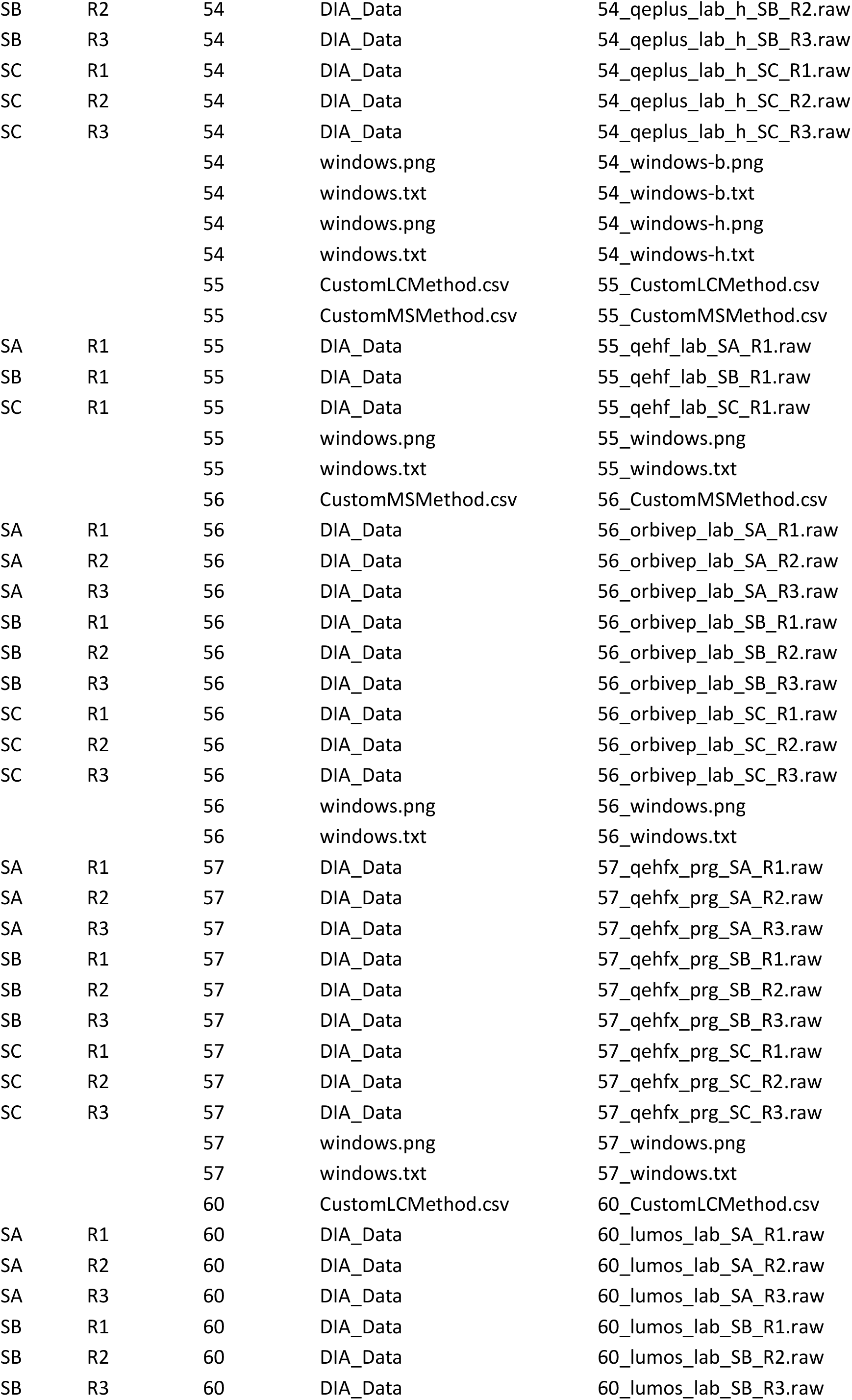
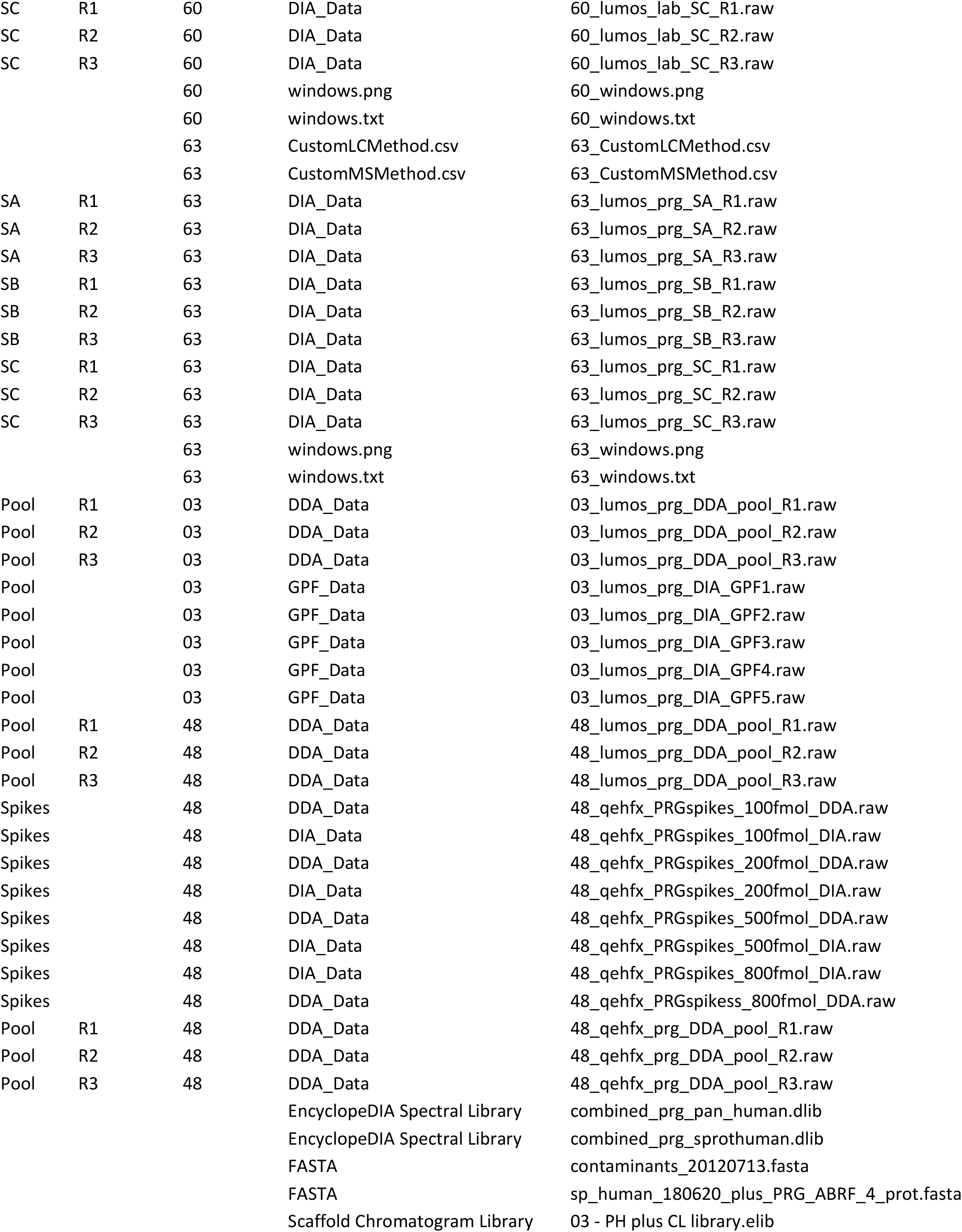
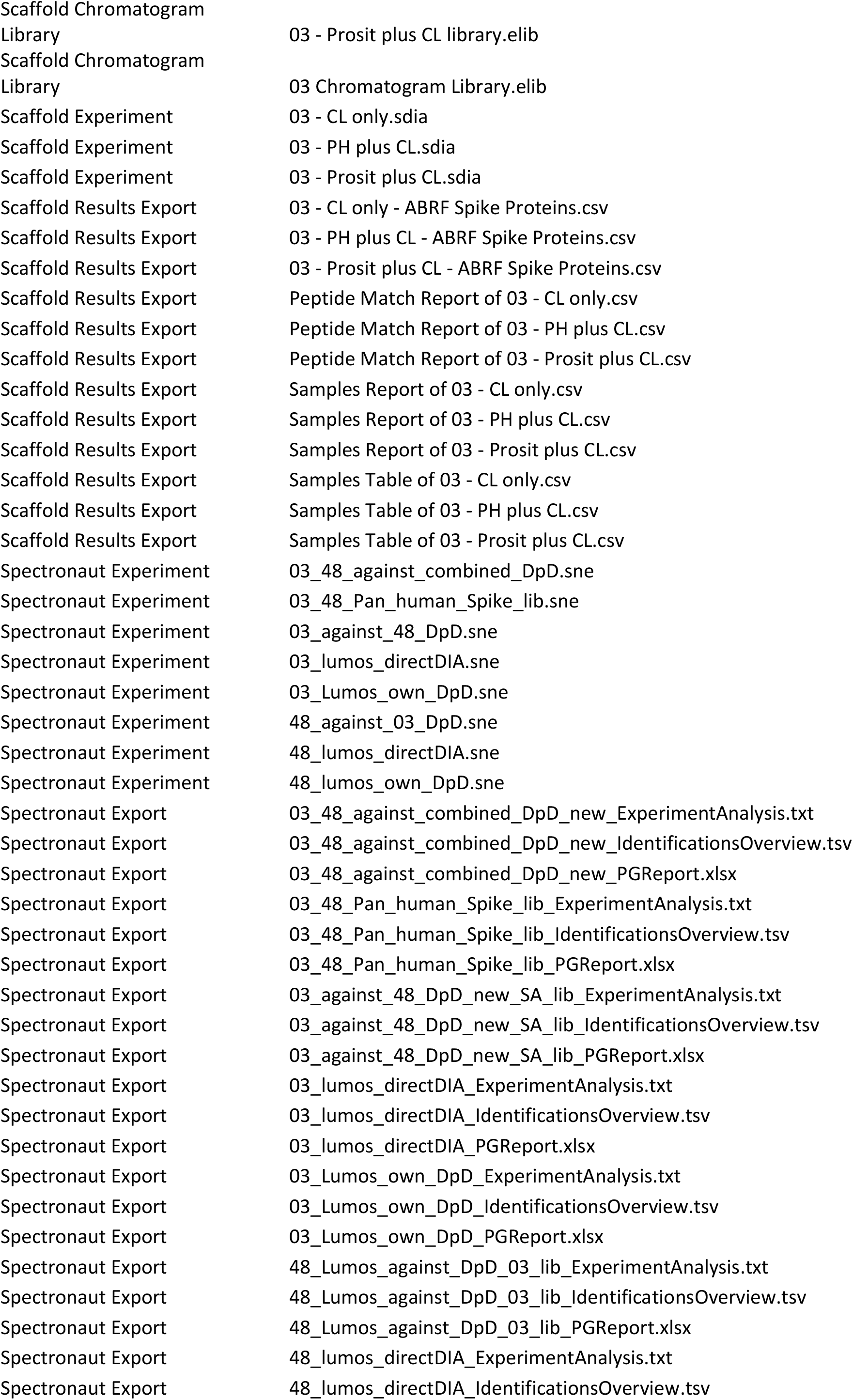
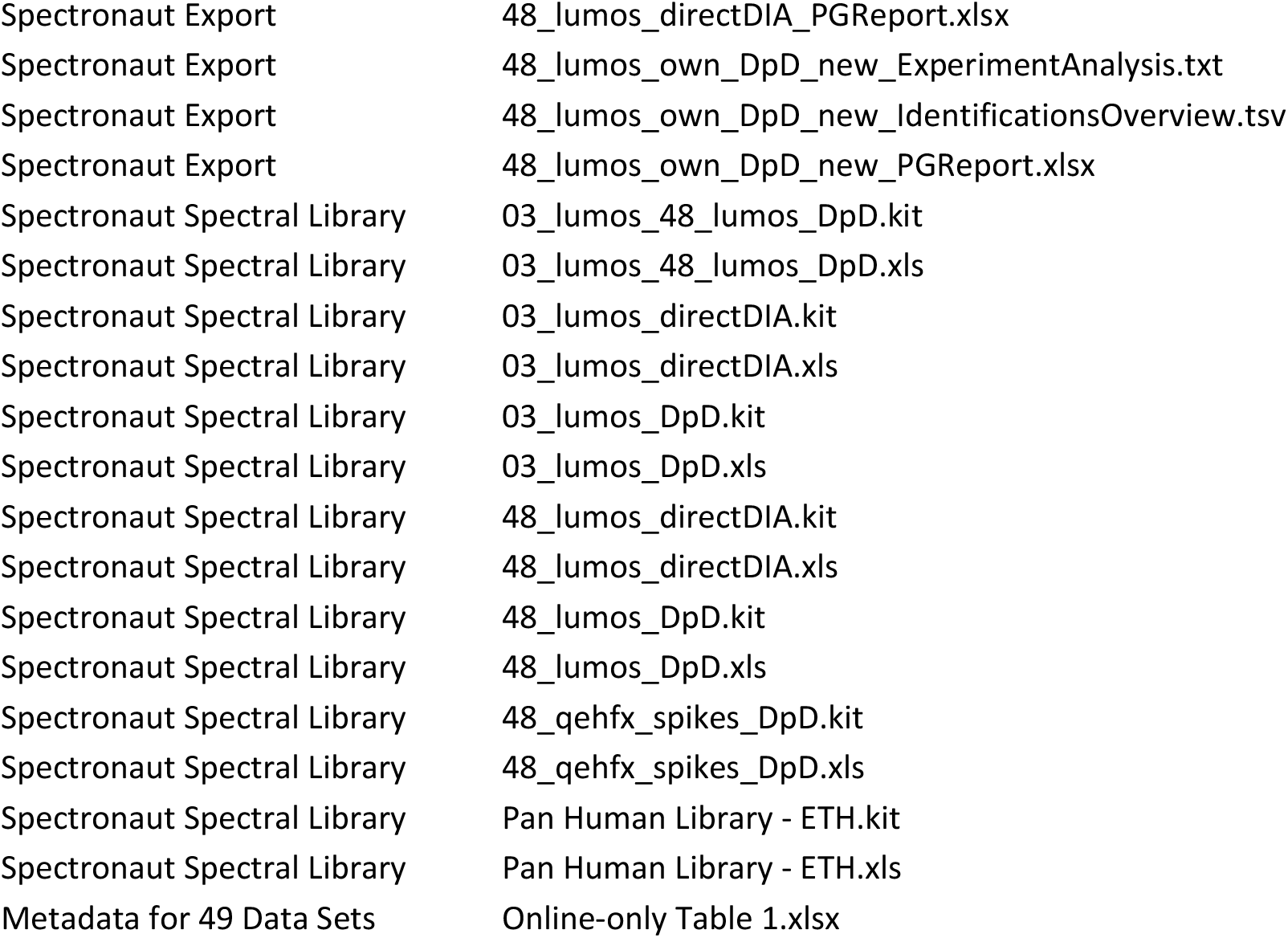

| Sample | Replicate | Participant | Data Type | Data |
| --- | --- | --- | --- | --- |
| SA | R1 | 01 | DIA_Data | 01_qehfx_lab_SA_R1.raw |
| SA | R2 | 01 | DIA_Data | 01_qehfx_lab_SA_R2.raw |
| SA | R3 | 01 | DIA_Data | 01_qehfx_lab_SA_R3.raw |
| SB | R1 | 01 | DIA_Data | 01_qehfx_lab_SB_R1.raw |
| SB | R2 | 01 | DIA_Data | 01_qehfx_lab_SB_R2.raw |
| SB | R3 | 01 | DIA_Data | 01_qehfx_lab_SB_R3.raw |
| SC | R1 | 01 | DIA_Data | 01_qehfx_lab_SC_R1.raw |
| SC | R2 | 01 | DIA_Data | 01_qehfx_lab_SC_R2.raw |
| SC | R3 | 01 | DIA_Data | 01_qehfx_lab_SC_R3.raw |
|  |  | 01 | windows.png | 01_windows.png |
|  |  | 01 | windows.txt | 01_windows.txt |
|  |  | 03 | instrument_method | 03_lumos_156min_15kMS2_62_14mz.meth |
| SA | R1 | 03 | DIA_Data | 03_lumos_prg_SA_R1.raw |
| SA | R2 | 03 | DIA_Data | 03_lumos_prg_SA_R2.raw |
| SA | R3 | 03 | DIA_Data | 03_lumos_prg_SA_R3.raw |
| SB | R1 | 03 | DIA_Data | 03_lumos_prg_SB_R1.raw |
| SB | R2 | 03 | DIA_Data | 03_lumos_prg_SB_R2.raw |
| SB | R3 | 03 | DIA_Data | 03_lumos_prg_SB_R3.raw |
| SC | R1 | 03 | DIA_Data | 03_lumos_prg_SC_R1.raw |
| SC | R2 | 03 | DIA_Data | 03_lumos_prg_SC_R2.raw |
| SC | R3 | 03 | DIA_Data | 03_lumos_prg_SC_R3.raw |
|  |  | 03 | windows.png | 03_windows.png |
|  |  | 03 | windows.txt | 03_windows.txt |
|  |  | 04 | CustomMSMethod.csv | 04_CustomMSMethod.csv |
| SA | R1 | 04 | DIA_Data | 04_orbive_lab_SA_R1.raw |
| SA | R2 | 04 | DIA_Data | 04_orbive_lab_SA_R2.raw |
| SA | R3 | 04 | DIA_Data | 04_orbive_lab_SA_R3.raw |
| SB | R1 | 04 | DIA_Data | 04_orbive_lab_SB_R1.raw |
| SB | R2 | 04 | DIA_Data | 04_orbive_lab_SB_R2.raw |
| SB | R3 | 04 | DIA_Data | 04_orbive_lab_SB_R3.raw |
| SC | R1 | 04 | DIA_Data | 04_orbive_lab_SC_R1.raw |
| SC | R2 | 04 | DIA_Data | 04_orbive_lab_SC_R2.raw |
| SC | R3 | 04 | DIA_Data | 04_orbive_lab_SC_R3.raw |
|  |  | 04 | windows.png | 04_windows.png |
|  |  | 04 | windows.txt | 04_windows.txt |
|  |  | 05 | CustomLCMethod.csv | 05_CustomLCMethod.csv |
|  |  | 05 | CustomMSMethod.csv | 05_CustomMSMethod.csv |
| SA | R1 | 05 | DIA_Data | 05_lumos_lab_SA_R1.raw |
| SA | R2 | 05 | DIA_Data | 05_lumos_lab_SA_R2.raw |
| SA | R3 | 05 | DIA_Data | 05_lumos_lab_SA_R3.raw |
| SB | R1 | 05 | DIA_Data | 05_lumos_lab_SB_R1.raw |
| SB | R2 | 05 | DIA_Data | 05_lumos_lab_SB_R2.raw |
| SB | R3 | 05 | DIA_Data | 05_lumos_lab_SB_R3.raw |
| SC | R1 | 05 | DIA_Data | 05_lumos_lab_SC_R1.raw |
| SC | R2 | 05 | DIA_Data | 05_lumos_lab_SC_R2.raw |
| SC | R3 | 05 | DIA_Data | 05_lumos_lab_SC_R3.raw |
|  |  | 05 | windows.png | 05_windows.png |
|  |  | 05 | windows.txt | 05_windows.txt |
| SA | R1 | 06 | DIA_Data | 06_lumos_lab_SA_R1.raw |
| SA | R2 | 06 | DIA_Data | 06_lumos_lab_SA_R2.raw |
| SA | R3 | 06 | DIA_Data | 06_lumos_lab_SA_R3.raw |
| SB | R1 | 06 | DIA_Data | 06_lumos_lab_SB_R1.raw |
| SB | R2 | 06 | DIA_Data | 06_lumos_lab_SB_R2.raw |
| SB | R3 | 06 | DIA_Data | 06_lumos_lab_SB_R3.raw |
| SC | R1 | 06 | DIA_Data | 06_lumos_lab_SC_R1.raw |
| SC | R2 | 06 | DIA_Data | 06_lumos_lab_SC_R2.raw |
| SC | R3 | 06 | DIA_Data | 06_lumos_lab_SC_R3.raw |
|  |  | 06 | 06_windows.png | 06_windows.png |
|  |  | 06 | 06_windows.txt | 06_windows.txt |
| SA | R1 | 07 | DIA_Data | 07_fusion_prg15k_SA_R1.raw |
| SA | R2 | 07 | DIA_Data | 07_fusion_prg15k_SA_R2.raw |
| SA | R3 | 07 | DIA_Data | 07_fusion_prg15k_SA_R3.raw |
| SB | R1 | 07 | DIA_Data | 07_fusion_prg15k_SB_R1.raw |
| SB | R2 | 07 | DIA_Data | 07_fusion_prg15k_SB_R2.raw |
| SB | R3 | 07 | DIA_Data | 07_fusion_prg15k_SB_R3.raw |
| SC | R1 | 07 | DIA_Data | 07_fusion_prg15k_SC_R1.raw |
| SC | R2 | 07 | DIA_Data | 07_fusion_prg15k_SC_R2.raw |
| SC | R3 | 07 | DIA_Data | 07_fusion_prg15k_SC_R3.raw |
| SA | R1 | 07 | DIA_Data | 07_fusion_prg30k_SA_R1.raw |
| SA | R3 | 07 | DIA_Data | 07_fusion_prg30k_SA_R3.raw |
| SB | R2 | 07 | DIA_Data | 07_fusion_prg30k_SB_R2.raw |
| SB | R3 | 07 | DIA_Data | 07_fusion_prg30k_SB_R3.raw |
| SC | R1 | 07 | DIA_Data | 07_fusion_prg30k_SC_R1.raw |
| SC | R2 | 07 | DIA_Data | 07_fusion_prg30k_SC_R2.raw |
| SC | R3 | 07 | DIA_Data | 07_fusion_prg30k_SC_R3.raw |
|  |  | 07 | DIA_Data | 07_windows_15k.png |
|  |  | 07 | DIA_Data | 07_windows_15k.txt |
|  |  | 07 | DIA_Data | 07_windows_30k.png |
|  |  | 07 | DIA_Data | 07_windows_30k.txt |
| SA | R1 | 08 | DIA_Data | 08_fusion_prg_SA_R1.raw |
| SA | R2 | 08 | DIA_Data | 08_fusion_prg_SA_R2.raw |
| SA | R3 | 08 | DIA_Data | 08_fusion_prg_SA_R3.raw |
| SB | R1 | 08 | DIA_Data | 08_fusion_prg_SB_R1.raw |
| SB | R2 | 08 | DIA_Data | 08_fusion_prg_SB_R2.raw |
| SB | R3 | 08 | DIA_Data | 08_fusion_prg_SB_R3.raw |
| SC | R1 | 08 | DIA_Data | 08_fusion_prg_SC_R1.raw |
| SC | R2 | 08 | DIA_Data | 08_fusion_prg_SC_R2.raw |
| SC | R3 | 08 | DIA_Data | 08_fusion_prg_SC_R3.raw |
|  |  | 08 | windows.png | 08_windows.png |
|  |  | 08 | windows.txt | 08_windows.txt |
| SA | R1 | 10 | DIA_Data | 10_qehf_prg_SA_R1.raw |
| SA | R2 | 10 | DIA_Data | 10_qehf_prg_SA_R2.raw |
| SA | R3 | 10 | DIA_Data | 10_qehf_prg_SA_R3.raw |
| SB | R1 | 10 | DIA_Data | 10_qehf_prg_SB_R1.raw |
| SB | R2 | 10 | DIA_Data | 10_qehf_prg_SB_R2.raw |
| SB | R3 | 10 | DIA_Data | 10_qehf_prg_SB_R3.raw |
| SC | R1 | 10 | DIA_Data | 10_qehf_prg_SC_R1.raw |
| SC | R2 | 10 | DIA_Data | 10_qehf_prg_SC_R2.raw |
| SC | R3 | 10 | DIA_Data | 10_qehf_prg_SC_R3.raw |
|  |  | 10 | windows.png | 10_windows.png |
|  |  | 10 | windows.txt | 10_windows.txt |
| SA | R1 | 13 | DIA_Data | 13_fusion_prg_SA_R1.raw |
| SA | R2 | 13 | DIA_Data | 13_fusion_prg_SA_R2.raw |
| SA | R3 | 13 | DIA_Data | 13_fusion_prg_SA_R3.raw |
| SB | R1 | 13 | DIA_Data | 13_fusion_prg_SB_R1.raw |
| SB | R2 | 13 | DIA_Data | 13_fusion_prg_SB_R2.raw |
| SB | R3 | 13 | DIA_Data | 13_fusion_prg_SB_R3.raw |
| SC | R1 | 13 | DIA_Data | 13_fusion_prg_SC_R1.raw |
| SC | R2 | 13 | DIA_Data | 13_fusion_prg_SC_R2.raw |
| SC | R3 | 13 | DIA_Data | 13_fusion_prg_SC_R3.raw |
|  |  | 13 | windows.png | 13_windows.png |
|  |  | 13 | windows.txt | 13_windows.txt |
|  |  | 14 | CustomLCMethod.csv | 14_CustomLCMethod.csv |
|  |  | 14 | CustomMSMethod.csv | 14_CustomMSMethod.csv |
| SA | R1 | 14 | DIA_Data | 14_tt6600_lab_SA_R1.wiff |
| SA | R1 | 14 | DIA_Data | 14_tt6600_lab_SA_R1.wiff.scan |
| SA | R2 | 14 | DIA_Data | 14_tt6600_lab_SA_R2.wiff |
| SA | R2 | 14 | DIA_Data | 14_tt6600_lab_SA_R2.wiff.scan |
| SA | R3 | 14 | DIA_Data | 14_tt6600_lab_SA_R3.wiff |
| SA | R3 | 14 | DIA_Data | 14_tt6600_lab_SA_R3.wiff.scan |
| SB | R1 | 14 | DIA_Data | 14_tt6600_lab_SB_R1.wiff |
| SB | R1 | 14 | DIA_Data | 14_tt6600_lab_SB_R1.wiff.scan |
| SB | R2 | 14 | DIA_Data | 14_tt6600_lab_SB_R2.wiff |
| SB | R2 | 14 | DIA_Data | 14_tt6600_lab_SB_R2.wiff.scan |
| SB | R3 | 14 | DIA_Data | 14_tt6600_lab_SB_R3.wiff |
| SB | R3 | 14 | DIA_Data | 14_tt6600_lab_SB_R3.wiff.scan |
| SC | R1 | 14 | DIA_Data | 14_tt6600_lab_SC_R1.wiff |
| SC | R1 | 14 | DIA_Data | 14_tt6600_lab_SC_R1.wiff.scan |
| SC | R2 | 14 | DIA_Data | 14_tt6600_lab_SC_R2.wiff |
| SC | R2 | 14 | DIA_Data | 14_tt6600_lab_SC_R2.wiff.scan |
| SC | R3 | 14 | DIA_Data | 14_tt6600_lab_SC_R3.wiff |
| SC | R3 | 14 | DIA_Data | 14_tt6600_lab_SC_R3.wiff.scan |
|  |  | 14 | windows.png | 14_windows.png |
|  |  | 14 | windows.txt | 14_windows.txt |
|  |  | 15 | CustomLCMethod.csv | 15_CustomLCMethod.csv |
|  |  | 15 | CustomMSMethod.csv | 15_CustomMSMethod.csv |
| SA | R1 | 15 | DIA_Data | 15_orbielite_lab_SA_R1.raw |
| SA | R2 | 15 | DIA_Data | 15_orbielite_lab_SA_R2.raw |
| SA | R3 | 15 | DIA_Data | 15_orbielite_lab_SA_R3.raw |
| SC | R1 | 15 | DIA_Data | 15_orbielite_lab_SC_R1.raw |
| SC | R2 | 15 | DIA_Data | 15_orbielite_lab_SC_R2.raw |
| SC | R3 | 15 | DIA_Data | 15_orbielite_lab_SC_R3.raw |
|  |  | 15 | windows.png | 15_windows.png |
|  |  | 15 | windows.txt | 15_windows.txt |
| SA | R1 | 16 | DIA_Data | 16_tt6600_prg_SA_R1.wiff |
| SA | R1 | 16 | DIA_Data | 16_tt6600_prg_SA_R1.wiff.scan |
| SA | R2 | 16 | DIA_Data | 16_tt6600_prg_SA_R2.wiff |
| SA | R2 | 16 | DIA_Data | 16_tt6600_prg_SA_R2.wiff.scan |
| SA | R3 | 16 | DIA_Data | 16_tt6600_prg_SA_R3.wiff |
| SA | R3 | 16 | DIA_Data | 16_tt6600_prg_SA_R3.wiff.scan |
| SB | R1 | 16 | DIA_Data | 16_tt6600_prg_SB_R1.wiff |
| SB | R1 | 16 | DIA_Data | 16_tt6600_prg_SB_R1.wiff.scan |
| SB | R2 | 16 | DIA_Data | 16_tt6600_prg_SB_R2.wiff |
| SB | R2 | 16 | DIA_Data | 16_tt6600_prg_SB_R2.wiff.scan |
| SB | R3 | 16 | DIA_Data | 16_tt6600_prg_SB_R3.wiff |
| SB | R3 | 16 | DIA_Data | 16_tt6600_prg_SB_R3.wiff.scan |
| SC | R1 | 16 | DIA_Data | 16_tt6600_prg_SC_R1.wiff |
| SC | R1 | 16 | DIA_Data | 16_tt6600_prg_SC_R1.wiff.scan |
| SC | R2 | 16 | DIA_Data | 16_tt6600_prg_SC_R2.wiff |
| SC | R2 | 16 | DIA_Data | 16_tt6600_prg_SC_R2.wiff.scan |
| SC | R3 | 16 | DIA_Data | 16_tt6600_prg_SC_R3.wiff |
| SC | R3 | 16 | DIA_Data | 16_tt6600_prg_SC_R3.wiff.scan |
|  |  | 16 | windows.png | 16_windows.png |
|  |  | 16 | windows.txt | 16_windows.txt |
| SA | R1 | 17 | DIA_Data | 17_qehfx_prg_SA_R1.raw |
| SA | R2 | 17 | DIA_Data | 17_qehfx_prg_SA_R2.raw |
| SA | R3 | 17 | DIA_Data | 17_qehfx_prg_SA_R3.raw |
| SB | R1 | 17 | DIA_Data | 17_qehfx_prg_SB_R1.raw |
| SB | R2 | 17 | DIA_Data | 17_qehfx_prg_SB_R2.raw |
| SB | R3 | 17 | DIA_Data | 17_qehfx_prg_SB_R3.raw |
| SC | R1 | 17 | DIA_Data | 17_qehfx_prg_SC_R1.raw |
| SC | R2 | 17 | DIA_Data | 17_qehfx_prg_SC_R2.raw |
| SC | R3 | 17 | DIA_Data | 17_qehfx_prg_SC_R3.raw |
|  |  | 17 | windows.png | 17_windows.png |
|  |  | 17 | windows.txt | 17_windows.txt |
|  |  | 18 | CustomLCMethod.csv | 18_CustomLCMethod.csv |
| SA | R1 | 18 | DIA_Data | 18_fusion_prg_SA_R1.raw |
| SA | R2 | 18 | DIA_Data | 18_fusion_prg_SA_R2.raw |
| SA | R3 | 18 | DIA_Data | 18_fusion_prg_SA_R3.raw |
| SB | R1 | 18 | DIA_Data | 18_fusion_prg_SB_R1.raw |
| SB | R2 | 18 | DIA_Data | 18_fusion_prg_SB_R2.raw |
| SB | R3 | 18 | DIA_Data | 18_fusion_prg_SB_R3.raw |
| SC | R1 | 18 | DIA_Data | 18_fusion_prg_SC_R1.raw |
| SC | R2 | 18 | DIA_Data | 18_fusion_prg_SC_R2.raw |
| SC | R3 | 18 | DIA_Data | 18_fusion_prg_SC_R3.raw |
|  |  | 18 | windows.png | 18_windows.png |
|  |  | 18 | windows.txt | 18_windows.txt |
|  |  | 19 | CustomLCMethod.csv | 19_CustomLCMethod.csv |
|  |  | 19 | CustomMSMethod.csv | 19_CustomMSMethod.csv |
| SA | R2 | 19 | DIA_Data | 19_qehfx_lab_SA_R2.raw |
| SA | R3 | 19 | DIA_Data | 19_qehfx_lab_SA_R3.raw |
| SB | R1 | 19 | DIA_Data | 19_qehfx_lab_SB_R1.raw |
| SB | R2 | 19 | DIA_Data | 19_qehfx_lab_SB_R2.raw |
| SB | R3 | 19 | DIA_Data | 19_qehfx_lab_SB_R3.raw |
| SC | R1 | 19 | DIA_Data | 19_qehfx_lab_SC_R1.raw |
| SC | R2 | 19 | DIA_Data | 19_qehfx_lab_SC_R2.raw |
| SC | R3 | 19 | DIA_Data | 19_qehfx_lab_SC_R3.raw |
|  |  | 19 | windows.png | 19_windows.png |
|  |  | 19 | windows.txt | 19_windows.txt |
|  |  | 20 | CustomLCmethod.csv | 20_CustomLCmethod.csv |
|  |  | 20 | CustomMSmethod.csv | 20_CustomMSmethod.csv |
|  |  | 20 | instrument_method | 20_PRG2018_SWATH.dam |
| SA | R1 | 20 | DIA_Data | 20_tt5600_lab_SA_R1.wiff |
| SA | R1 | 20 | DIA_Data | 20_tt5600_lab_SA_R1.wiff.scan |
| SA | R2 | 20 | DIA_Data | 20_tt5600_lab_SA_R2.wiff |
| SA | R2 | 20 | DIA_Data | 20_tt5600_lab_SA_R2.wiff.scan |
| SA | R3 | 20 | DIA_Data | 20_tt5600_lab_SA_R3.wiff |
| SA | R3 | 20 | DIA_Data | 20_tt5600_lab_SA_R3.wiff.scan |
| SB | R1 | 20 | DIA_Data | 20_tt5600_lab_SB_R1.wiff |
| SB | R1 | 20 | DIA_Data | 20_tt5600_lab_SB_R1.wiff.scan |
| SB | R2 | 20 | DIA_Data | 20_tt5600_lab_SB_R2.wiff |
| SB | R2 | 20 | DIA_Data | 20_tt5600_lab_SB_R2.wiff.scan |
| SB | R3 | 20 | DIA_Data | 20_tt5600_lab_SB_R3.wiff |
| SB | R3 | 20 | DIA_Data | 20_tt5600_lab_SB_R3.wiff.scan |
| SC | R1 | 20 | DIA_Data | 20_tt5600_lab_SC_R1.wiff |
| SC | R1 | 20 | DIA_Data | 20_tt5600_lab_SC_R1.wiff.scan |
| SC | R2 | 20 | DIA_Data | 20_tt5600_lab_SC_R2.wiff |
| SC | R2 | 20 | DIA_Data | 20_tt5600_lab_SC_R2.wiff.scan |
| SC | R3 | 20 | DIA_Data | 20_tt5600_lab_SC_R3.wiff |
| SC | R3 | 20 | DIA_Data | 20_tt5600_lab_SC_R3.wiff.scan |
|  |  | 20 | windows.png | 20_windows.png |
|  |  | 20 | windows.txt | 20_windows.txt |
|  |  | 21 | CustomLCMethod.csv | 21_CustomLCMethod.csv |
| SA | R1 | 21 | DIA_Data | 21_fusion_prg_SA_R1.raw |
| SA | R2 | 21 | DIA_Data | 21_fusion_prg_SA_R2.raw |
| SA | R3 | 21 | DIA_Data | 21_fusion_prg_SA_R3.raw |
| SB | R1 | 21 | DIA_Data | 21_fusion_prg_SB_R1.raw |
| SB | R2 | 21 | DIA_Data | 21_fusion_prg_SB_R2.raw |
| SB | R3 | 21 | DIA_Data | 21_fusion_prg_SB_R3.raw |
| SC | R1 | 21 | DIA_Data | 21_fusion_prg_SC_R1.raw |
| SC | R2 | 21 | DIA_Data | 21_fusion_prg_SC_R2.raw |
| SC | R3 | 21 | DIA_Data | 21_fusion_prg_SC_R3.raw |
|  |  | 21 | windows.png | 21_windows.png |
|  |  | 21 | windows.txt | 21_windows.txt |
|  |  | 23 | CustomLCMethod.csv | 23_CustomLCMethod.csv |
| SA | R1 | 23 | DIA_Data | 23_xevo_prg_SA_R1.raw |
| SA | R2 | 23 | DIA_Data | 23_xevo_prg_SA_R2.raw |
| SA | R3 | 23 | DIA_Data | 23_xevo_prg_SA_R3.raw |
| SB | R1 | 23 | DIA_Data | 23_xevo_prg_SB_R1.raw |
| SB | R2 | 23 | DIA_Data | 23_xevo_prg_SB_R2.raw |
| SB | R3 | 23 | DIA_Data | 23_xevo_prg_SB_R3.raw |
| SC | R1 | 23 | DIA_Data | 23_xevo_prg_SC_R1.raw |
| SC | R2 | 23 | DIA_Data | 23_xevo_prg_SC_R2.raw |
| SC | R3 | 23 | DIA_Data | 23_xevo_prg_SC_R3.raw |
| SA | R2 | 25 | DIA_Data | 25_fusion_prg_SA_R2.raw |
| SA | R3 | 25 | DIA_Data | 25_fusion_prg_SA_R3.raw |
| SB | R1 | 25 | DIA_Data | 25_fusion_prg_SB_R1.raw |
| SB | R2 | 25 | DIA_Data | 25_fusion_prg_SB_R2.raw |
| SB | R3 | 25 | DIA_Data | 25_fusion_prg_SB_R3.raw |
| SC | R1 | 25 | DIA_Data | 25_fusion_prg_SC_R1.raw |
| SC | R2 | 25 | DIA_Data | 25_fusion_prg_SC_R2.raw |
| SC | R3 | 25 | DIA_Data | 25_fusion_prg_SC_R3.raw |
|  |  | 25 | windows.png | 25_windows.png |
|  |  | 25 | windows.txt | 25_windows.txt |
|  |  | 26 | CustomLCMethod.csv | 26_CustomLCMethod.csv |
|  |  | 26 | CustomMSMethod.csv | 26_CustomMSMethod.csv |
| SA | R1 | 26 | DIA_Data | 26_tt5600_lab_SA_R1.wiff |
| SA | R1 | 26 | DIA_Data | 26_tt5600_lab_SA_R1.wiff.scan |
| SA | R2 | 26 | DIA_Data | 26_tt5600_lab_SA_R2.wiff |
| SA | R2 | 26 | DIA_Data | 26_tt5600_lab_SA_R2.wiff.scan |
| SA | R3 | 26 | DIA_Data | 26_tt5600_lab_SA_R3.wiff |
| SA | R3 | 26 | DIA_Data | 26_tt5600_lab_SA_R3.wiff.scan |
| SB | R1 | 26 | DIA_Data | 26_tt5600_lab_SB_R1.wiff |
| SB | R1 | 26 | DIA_Data | 26_tt5600_lab_SB_R1.wiff.scan |
| SB | R2 | 26 | DIA_Data | 26_tt5600_lab_SB_R2.wiff |
| SB | R2 | 26 | DIA_Data | 26_tt5600_lab_SB_R2.wiff.scan |
| SB | R3 | 26 | DIA_Data | 26_tt5600_lab_SB_R3.wiff |
| SB | R3 | 26 | DIA_Data | 26_tt5600_lab_SB_R3.wiff.scan |
| SC | R1 | 26 | DIA_Data | 26_tt5600_lab_SC_R1.wiff |
| SC | R1 | 26 | DIA_Data | 26_tt5600_lab_SC_R1.wiff.scan |
| SC | R2 | 26 | DIA_Data | 26_tt5600_lab_SC_R2.wiff |
| SC | R2 | 26 | DIA_Data | 26_tt5600_lab_SC_R2.wiff.scan |
| SC | R3 | 26 | DIA_Data | 26_tt5600_lab_SC_R3.wiff |
| SC | R3 | 26 | DIA_Data | 26_tt5600_lab_SC_R3.wiff.scan |
|  |  | 26 | windows.png | 26_windows.png |
|  |  | 26 | windows.txt | 26_windows.txt |
| SA | R1 | 28 | DIA_Data | 28_qehf_prg_SA_R1.raw |
| SA | R2 | 28 | DIA_Data | 28_qehf_prg_SA_R2.raw |
| SA | R3 | 28 | DIA_Data | 28_qehf_prg_SA_R3.raw |
| SB | R1 | 28 | DIA_Data | 28_qehf_prg_SB_R1.raw |
| SB | R2 | 28 | DIA_Data | 28_qehf_prg_SB_R2.raw |
| SB | R3 | 28 | DIA_Data | 28_qehf_prg_SB_R3.raw |
| SC | R1 | 28 | DIA_Data | 28_qehf_prg_SC_R1.raw |
| SC | R2 | 28 | DIA_Data | 28_qehf_prg_SC_R2.raw |
| SC | R3 | 28 | DIA_Data | 28_qehf_prg_SC_R3.raw |
|  |  | 28 | windows.png | 28_windows.png |
|  |  | 28 | windows.txt | 28_windows.txt |
| SA | R1 | 29 | DIA_Data | 29_lumos_SA_R1.raw |
| SA | R2 | 29 | DIA_Data | 29_lumos_SA_R2.raw |
| SA | R3 | 29 | DIA_Data | 29_lumos_SA_R3.raw |
| SB | R1 | 29 | DIA_Data | 29_lumos_SB_R1.raw |
| SB | R2 | 29 | DIA_Data | 29_lumos_SB_R2.raw |
| SB | R3 | 29 | DIA_Data | 29_lumos_SB_R3.raw |
| SC | R1 | 29 | DIA_Data | 29_lumos_SC_R1.raw |
| SC | R2 | 29 | DIA_Data | 29_lumos_SC_R2.raw |
| SC | R3 | 29 | DIA_Data | 29_lumos_SC_R3.raw |
|  |  | 29 | windows.png | 29_windows.png |
|  |  | 29 | windows.txt | 29_windows.txt |
| SA | R1 | 31 | DIA_Data | 31_lumos_prg_SA_R1.raw |
| SA | R2 | 31 | DIA_Data | 31_lumos_prg_SA_R2.raw |
| SA | R3 | 31 | DIA_Data | 31_lumos_prg_SA_R3.raw |
| SB | R1 | 31 | DIA_Data | 31_lumos_prg_SB_R1.raw |
| SB | R2 | 31 | DIA_Data | 31_lumos_prg_SB_R2.raw |
| SB | R3 | 31 | DIA_Data | 31_lumos_prg_SB_R3.raw |
| SC | R1 | 31 | DIA_Data | 31_lumos_prg_SC_R1.raw |
| SC | R2 | 31 | DIA_Data | 31_lumos_prg_SC_R2.raw |
| SC | R3 | 31 | DIA_Data | 31_lumos_prg_SC_R3.raw |
|  |  | 31 | windows.png | 31_windows.png |
|  |  | 31 | windows.txt | 31_windows.txt |
|  |  | 32 | CustomLCMethod.csv | 32_CustomLCMethod.csv |
|  |  | 32 | CustomMSMethod.csv | 32_CustomMSMethod.csv |
| SA | R1 | 32 | DIA_Data | 32_lumos_lab_SA_R1.raw |
| SA | R2 | 32 | DIA_Data | 32_lumos_lab_SA_R2.raw |
| SA | R3 | 32 | DIA_Data | 32_lumos_lab_SA_R3.raw |
| SB | R1 | 32 | DIA_Data | 32_lumos_lab_SB_R1.raw |
| SB | R2 | 32 | DIA_Data | 32_lumos_lab_SB_R2.raw |
| SB | R3 | 32 | DIA_Data | 32_lumos_lab_SB_R3.raw |
| SC | R1 | 32 | DIA_Data | 32_lumos_lab_SC_R1.raw |
| SC | R2 | 32 | DIA_Data | 32_lumos_lab_SC_R2.raw |
| SC | R3 | 32 | DIA_Data | 32_lumos_lab_SC_R3.raw |
|  |  | 32 | windows.png | 32_windows.png |
|  |  | 32 | windows.txt | 32_windows.txt |
| SA | R1 | 33 | DIA_Data | 33_lumos_prg_SA_R1.raw |
| SA | R2 | 33 | DIA_Data | 33_lumos_prg_SA_R2.raw |
| SA | R3 | 33 | DIA_Data | 33_lumos_prg_SA_R3.raw |
| SB | R1 | 33 | DIA_Data | 33_lumos_prg_SB_R1.raw |
| SB | R3 | 33 | DIA_Data | 33_lumos_prg_SB_R3.raw |
| SC | R1 | 33 | DIA_Data | 33_lumos_prg_SC_R1.raw |
| SC | R2 | 33 | DIA_Data | 33_lumos_prg_SC_R2.raw |
| SC | R3 | 33 | DIA_Data | 33_lumos_prg_SC_R3.raw |
|  |  | 33 | windows.png | 33_windows.png |
|  |  | 33 | windows.txt | 33_windows.txt |
| SA | R1 | 34 | DIA_Data | 34_qehf_prg_SA_R1.raw |
| SA | R2 | 34 | DIA_Data | 34_qehf_prg_SA_R2.raw |
| SA | R3 | 34 | DIA_Data | 34_qehf_prg_SA_R3.raw |
| SB | R1 | 34 | DIA_Data | 34_qehf_prg_SB_R1.raw |
| SB | R2 | 34 | DIA_Data | 34_qehf_prg_SB_R2.raw |
| SB | R3 | 34 | DIA_Data | 34_qehf_prg_SB_R3.raw |
| SC | R1 | 34 | DIA_Data | 34_qehf_prg_SC_R1.raw |
| SC | R2 | 34 | DIA_Data | 34_qehf_prg_SC_R2.raw |
| SC | R3 | 34 | DIA_Data | 34_qehf_prg_SC_R3.raw |
|  |  | 34 | windows.png | 34_windows.png |
|  |  | 34 | windows.txt | 34_windows.txt |
| SA | R1 | 35 | DIA_Data | 35_qehf_prg_SA_R1.raw |
| SA | R2 | 35 | DIA_Data | 35_qehf_prg_SA_R2.raw |
| SA | R3 | 35 | DIA_Data | 35_qehf_prg_SA_R3.raw |
| SB | R1 | 35 | DIA_Data | 35_qehf_prg_SB_R1.raw |
| SB | R2 | 35 | DIA_Data | 35_qehf_prg_SB_R2.raw |
| SB | R3 | 35 | DIA_Data | 35_qehf_prg_SB_R3.raw |
| SC | R1 | 35 | DIA_Data | 35_qehf_prg_SC_R1.raw |
| SC | R2 | 35 | DIA_Data | 35_qehf_prg_SC_R2.raw |
| SC | R3 | 35 | DIA_Data | 35_qehf_prg_SC_R3.raw |
|  |  | 35 | windows.png | 35_windows.png |
|  |  | 35 | windows.txt | 35_windows.txt |
|  |  | 36 | CustomLCMethod.csv | 36_CustomLCMethod.csv |
|  |  | 36 | CustomMSMethod.csv | 36_CustomMSMethod.csv |
| SA | R1 | 36 | DIA_Data | 36_qehf_lab_SA_R1.raw |
| SA | R2 | 36 | DIA_Data | 36_qehf_lab_SA_R2.raw |
| SA | R3 | 36 | DIA_Data | 36_qehf_lab_SA_R3.raw |
| SB | R1 | 36 | DIA_Data | 36_qehf_lab_SB_R1.raw |
| SB | R2 | 36 | DIA_Data | 36_qehf_lab_SB_R2.raw |
| SB | R3 | 36 | DIA_Data | 36_qehf_lab_SB_R3.raw |
| SC | R1 | 36 | DIA_Data | 36_qehf_lab_SC_R1.raw |
| SC | R2 | 36 | DIA_Data | 36_qehf_lab_SC_R2.raw |
| SC | R3 | 36 | DIA_Data | 36_qehf_lab_SC_R3.raw |
|  |  | 36 | windows.png | 36_windows.png |
|  |  | 36 | windows.txt | 36_windows.txt |
| SA | R1 | 37 | DIA_Data | 37_fusion_prg_SA_R1.raw |
| SA | R2 | 37 | DIA_Data | 37_fusion_prg_SA_R2.raw |
| SA | R3 | 37 | DIA_Data | 37_fusion_prg_SA_R3.raw |
| SB | R1 | 37 | DIA_Data | 37_fusion_prg_SB_R1.raw |
| SB | R2 | 37 | DIA_Data | 37_fusion_prg_SB_R2.raw |
| SB | R3 | 37 | DIA_Data | 37_fusion_prg_SB_R3.raw |
| SC | R1 | 37 | DIA_Data | 37_fusion_prg_SC_R1.raw |
| SC | R2 | 37 | DIA_Data | 37_fusion_prg_SC_R2.raw |
| SC | R3 | 37 | DIA_Data | 37_fusion_prg_SC_R3.raw |
|  |  | 37 | windows.png | 37_windows.png |
|  |  | 37 | windows.txt | 37_windows.txt |
|  |  | 39 | CustomLCMethod.csv | 39_CustomLCMethod.csv |
|  |  | 39 | CustomMSMethod.csv | 39_CustomMSMethod.csv |
| SA | R1 | 39 | DIA_Data | 39_tt5600_lab_SA_R1.wiff |
| SA | R1 | 39 | DIA_Data | 39_tt5600_lab_SA_R1.wiff.scan |
| SA | R2 | 39 | DIA_Data | 39_tt5600_lab_SA_R2.wiff |
| SA | R2 | 39 | DIA_Data | 39_tt5600_lab_SA_R2.wiff.scan |
| SA | R3 | 39 | DIA_Data | 39_tt5600_lab_SA_R3.wiff |
| SA | R3 | 39 | DIA_Data | 39_tt5600_lab_SA_R3.wiff.scan |
| SB | R1 | 39 | DIA_Data | 39_tt5600_lab_SB_R1.wiff |
| SB | R1 | 39 | DIA_Data | 39_tt5600_lab_SB_R1.wiff.scan |
| SB | R2 | 39 | DIA_Data | 39_tt5600_lab_SB_R2.wiff |
| SB | R2 | 39 | DIA_Data | 39_tt5600_lab_SB_R2.wiff.scan |
| SB | R3 | 39 | DIA_Data | 39_tt5600_lab_SB_R3.wiff |
| SB | R3 | 39 | DIA_Data | 39_tt5600_lab_SB_R3.wiff.scan |
| SC | R1 | 39 | DIA_Data | 39_tt5600_lab_SC_R1.wiff |
| SC | R1 | 39 | DIA_Data | 39_tt5600_lab_SC_R1.wiff.scan |
| SC | R2 | 39 | DIA_Data | 39_tt5600_lab_SC_R2.wiff |
| SC | R2 | 39 | DIA_Data | 39_tt5600_lab_SC_R2.wiff.scan |
| SC | R3 | 39 | DIA_Data | 39_tt5600_lab_SC_R3.wiff |
| SC | R3 | 39 | DIA_Data | 39_tt5600_lab_SC_R3.wiff.scan |
|  |  | 39 | windows.png | 39_windows.png |
|  |  | 39 | windows.txt | 39_windows.txt |
| SA | R1 | 44 | DIA_Data | 44_qehfx_prg_SA_R1.raw |
| SA | R2 | 44 | DIA_Data | 44_qehfx_prg_SA_R2.raw |
| SA | R3 | 44 | DIA_Data | 44_qehfx_prg_SA_R3.raw |
| SB | R1 | 44 | DIA_Data | 44_qehfx_prg_SB_R1.raw |
| SB | R2 | 44 | DIA_Data | 44_qehfx_prg_SB_R2.raw |
| SB | R3 | 44 | DIA_Data | 44_qehfx_prg_SB_R3.raw |
| SC | R1 | 44 | DIA_Data | 44_qehfx_prg_SC_R1.raw |
| SC | R2 | 44 | DIA_Data | 44_qehfx_prg_SC_R2.raw |
| SC | R3 | 44 | DIA_Data | 44_qehfx_prg_SC_R3.raw |
|  |  | 44 | windows.png | 44_windows.png |
|  |  | 44 | windows.txt | 44_windows.txt |
| SA | R1 | 45 | DIA_Data | 45_qeplus_prg_SA_R1.raw |
| SA | R2 | 45 | DIA_Data | 45_qeplus_prg_SA_R2.raw |
| SA | R3 | 45 | DIA_Data | 45_qeplus_prg_SA_R3.raw |
| SB | R1 | 45 | DIA_Data | 45_qeplus_prg_SB_R1.raw |
| SB | R2 | 45 | DIA_Data | 45_qeplus_prg_SB_R2.raw |
| SB | R3 | 45 | DIA_Data | 45_qeplus_prg_SB_R3.raw |
| SC | R1 | 45 | DIA_Data | 45_qeplus_prg_SC_R1.raw |
| SC | R2 | 45 | DIA_Data | 45_qeplus_prg_SC_R2.raw |
| SC | R3 | 45 | DIA_Data | 45_qeplus_prg_SC_R3.raw |
|  |  | 45 | windows.png | 45_windows.png |
|  |  | 45 | windows.txt | 45_windows.txt |
|  |  | 46 | CustomLCMethod.csv | 46_CustomLCMethod.csv |
|  |  | 46 | CustomMSMethod.csv | 46_CustomMSMethod.csv |
|  |  | 46 | windows.png | 46_lab_method_windows.png |
|  |  | 46 | windows.txt | 46_lab_method_windows.txt |
| SA | R1 | 46 | DIA_Data | 46_lumos_lab_SA_R1.raw |
| SA | R2 | 46 | DIA_Data | 46_lumos_lab_SA_R2.raw |
| SA | R3 | 46 | DIA_Data | 46_lumos_lab_SA_R3.raw |
| SB | R1 | 46 | DIA_Data | 46_lumos_lab_SB_R1.raw |
| SB | R2 | 46 | DIA_Data | 46_lumos_lab_SB_R2.raw |
| SB | R3 | 46 | DIA_Data | 46_lumos_lab_SB_R3.raw |
| SC | R1 | 46 | DIA_Data | 46_lumos_lab_SC_R1.raw |
| SC | R2 | 46 | DIA_Data | 46_lumos_lab_SC_R2.raw |
| SC | R3 | 46 | DIA_Data | 46_lumos_lab_SC_R3.raw |
| SA | R1 | 46 | DIA_Data | 46_lumos_prg_SA_R1.raw |
| SA | R2 | 46 | DIA_Data | 46_lumos_prg_SA_R2.raw |
| SA | R3 | 46 | DIA_Data | 46_lumos_prg_SA_R3.raw |
| SA | R4 | 46 | DIA_Data | 46_lumos_prg_SA_R4.raw |
| SB | R1 | 46 | DIA_Data | 46_lumos_prg_SB_R1.raw |
| SB | R2 | 46 | DIA_Data | 46_lumos_prg_SB_R2.raw |
| SB | R3 | 46 | DIA_Data | 46_lumos_prg_SB_R3.raw |
| SB | R4 | 46 | DIA_Data | 46_lumos_prg_SB_R4.raw |
| SC | R1 | 46 | DIA_Data | 46_lumos_prg_SC_R1.raw |
| SC | R2 | 46 | DIA_Data | 46_lumos_prg_SC_R2.raw |
| SC | R3 | 46 | DIA_Data | 46_lumos_prg_SC_R3.raw |
| SC | R4 | 46 | DIA_Data | 46_lumos_prg_SC_R4.raw |
|  |  | 46 | windows.png | 46_prg_method_windows.png |
|  |  | 46 | windows.txt | 46_prg_method_windows.txt |
|  |  | 47 | CustomLCMethod.csv | 47_CustomLCMethod.csv |
|  |  | 47 | CustomMSMethod.csv | 47_CustomMSMethod.csv |
|  |  | 47 | instrument_method | 47_DIA_PRG_25wind_24mz_400_1000.meth |
| SA | R1 | 47 | DIA_Data | 47_qe_lab_SA_R1.raw |
| SA | R2 | 47 | DIA_Data | 47_qe_lab_SA_R2.raw |
| SA | R3 | 47 | DIA_Data | 47_qe_lab_SA_R3.raw |
| SB | R1 | 47 | DIA_Data | 47_qe_lab_SB_R1.raw |
| SB | R2 | 47 | DIA_Data | 47_qe_lab_SB_R2.raw |
| SB | R3 | 47 | DIA_Data | 47_qe_lab_SB_R3.raw |
| SC | R1 | 47 | DIA_Data | 47_qe_lab_SC_R1.raw |
| SC | R2 | 47 | DIA_Data | 47_qe_lab_SC_R2.raw |
| SC | R3 | 47 | DIA_Data | 47_qe_lab_SC_R3.raw |
|  |  | 47 | windows.png | 47_windows.png |
|  |  | 47 | windows.txt | 47_windows.txt |
| SA | R1 | 48 | DIA_Data | 48_lumos_prg_SA_R1.raw |
| SA | R2 | 48 | DIA_Data | 48_lumos_prg_SA_R2.raw |
| SA | R3 | 48 | DIA_Data | 48_lumos_prg_SA_R3.raw |
| SB | R1 | 48 | DIA_Data | 48_lumos_prg_SB_R1.raw |
| SB | R2 | 48 | DIA_Data | 48_lumos_prg_SB_R2.raw |
| SB | R3 | 48 | DIA_Data | 48_lumos_prg_SB_R3.raw |
| SC | R1 | 48 | DIA_Data | 48_lumos_prg_SC_R1.raw |
| SC | R2 | 48 | DIA_Data | 48_lumos_prg_SC_R2.raw |
| SC | R3 | 48 | DIA_Data | 48_lumos_prg_SC_R3.raw |
| SA | R1 | 48 | DIA_Data | 48_qehfx_prg_SA_R1.raw |
| SA | R2 | 48 | DIA_Data | 48_qehfx_prg_SA_R2.raw |
| SA | R3 | 48 | DIA_Data | 48_qehfx_prg_SA_R3.raw |
| SB | R1 | 48 | DIA_Data | 48_qehfx_prg_SB_R1.raw |
| SB | R2 | 48 | DIA_Data | 48_qehfx_prg_SB_R2.raw |
| SB | R3 | 48 | DIA_Data | 48_qehfx_prg_SB_R3.raw |
| SC | R1 | 48 | DIA_Data | 48_qehfx_prg_SC_R1.raw |
| SC | R2 | 48 | DIA_Data | 48_qehfx_prg_SC_R2.raw |
| SC | R3 | 48 | DIA_Data | 48_qehfx_prg_SC_R3.raw |
|  |  | 48 | windows-lumos.png | 48_windows-lumos.png |
|  |  | 48 | windows-lumos.txt | 48_windows-lumos.txt |
|  |  | 48 | windows-qehfx.png | 48_windows-qehfx.png |
|  |  | 48 | windows-qehfx.txt | 48_windows-qehfx.txt |
|  |  | 49 | CustomMSMethod.csv | 49_CustomMSMethod.csv |
|  |  | 49 | CustomIsolationWindows.csv | 49_CustomIsolationWindows.csv |
| SA | R1 | 49 | DIA_Data | 49_qe_lab_SA_R1.raw |
| SA | R2 | 49 | DIA_Data | 49_qe_lab_SA_R2.raw |
| SA | R3 | 49 | DIA_Data | 49_qe_lab_SA_R3.raw |
| SB | R1 | 49 | DIA_Data | 49_qe_lab_SB_R1.raw |
| SB | R2 | 49 | DIA_Data | 49_qe_lab_SB_R2.raw |
| SB | R3 | 49 | DIA_Data | 49_qe_lab_SB_R3.raw |
| SC | R1 | 49 | DIA_Data | 49_qe_lab_SC_R1.raw |
| SC | R2 | 49 | DIA_Data | 49_qe_lab_SC_R2.raw |
| SC | R3 | 49 | DIA_Data | 49_qe_lab_SC_R3.raw |
|  |  | 49 | windows.png | 49_windows.png |
|  |  | 49 | windows.txt | 49_windows.txt |
|  |  | 50 | CustomLCMethod.csv | 50_CustomLCMethod.csv |
| SA | R1 | 50 | DIA_Data | 50_qehfx_prg_SA_R1.raw |
| SA | R2 | 50 | DIA_Data | 50_qehfx_prg_SA_R2.raw |
| SA | R3 | 50 | DIA_Data | 50_qehfx_prg_SA_R3.raw |
| SB | R1 | 50 | DIA_Data | 50_qehfx_prg_SB_R1.raw |
| SB | R2 | 50 | DIA_Data | 50_qehfx_prg_SB_R2.raw |
| SB | R3 | 50 | DIA_Data | 50_qehfx_prg_SB_R3.raw |
| SC | R1 | 50 | DIA_Data | 50_qehfx_prg_SC_R1.raw |
| SC | R2 | 50 | DIA_Data | 50_qehfx_prg_SC_R2.raw |
| SC | R3 | 50 | DIA_Data | 50_qehfx_prg_SC_R3.raw |
|  |  | 50 | windows.png | 50_windows.png |
|  |  | 50 | windows.txt | 50_windows.txt |
| SA | R1 | 52 | DIA_Data | 52_fusion_lab_SA_R1.raw |
| SA | R2 | 52 | DIA_Data | 52_fusion_lab_SA_R2.raw |
| SA | R3 | 52 | DIA_Data | 52_fusion_lab_SA_R3.raw |
| SB | R1 | 52 | DIA_Data | 52_fusion_lab_SB_R1.raw |
| SB | R2 | 52 | DIA_Data | 52_fusion_lab_SB_R2.raw |
| SB | R3 | 52 | DIA_Data | 52_fusion_lab_SB_R3.raw |
| SC | R1 | 52 | DIA_Data | 52_fusion_lab_SC_R1.raw |
| SC | R2 | 52 | DIA_Data | 52_fusion_lab_SC_R2.raw |
| SC | R3 | 52 | DIA_Data | 52_fusion_lab_SC_R3.raw |
|  |  | 52 | windows.png | 52_windows.png |
|  |  | 52 | windows.txt | 52_windows.txt |
| SA | R1 | 53 | DIA_Data | 53_qe_prg_SA_R1.raw |
| SA | R2 | 53 | DIA_Data | 53_qe_prg_SA_R2.raw |
| SA | R3 | 53 | DIA_Data | 53_qe_prg_SA_R3.raw |
| SB | R1 | 53 | DIA_Data | 53_qe_prg_SB_R1.raw |
| SB | R2 | 53 | DIA_Data | 53_qe_prg_SB_R2.raw |
| SB | R3 | 53 | DIA_Data | 53_qe_prg_SB_R3.raw |
| SC | R1 | 53 | DIA_Data | 53_qe_prg_SC_R1.raw |
| SC | R2 | 53 | DIA_Data | 53_qe_prg_SC_R2.raw |
| SC | R3 | 53 | DIA_Data | 53_qe_prg_SC_R3.raw |
|  |  | 53 | windows.png | 53_windows.png |
|  |  | 53 | windows.txt | 53_windows.txt |
|  |  | 54 | CustomLCMethod.csv | 54_CustomLCMethod_bandh.csv |
|  |  | 54 | CustomMSMethod.csv | 54_CustomMSMethod_b.csv |
|  |  | 54 | CustomMSMethod.csv | 54_CustomMSMethod_h.csv |
|  |  | 54 | instrument_method | 54_DIA_140min_b.meth |
|  |  | 54 | instrument_method | 54_DIA_140min_h.meth |
| SA | R1 | 54 | DIA_Data | 54_qeplus_lab_b_SA_R1.raw |
| SA | R2 | 54 | DIA_Data | 54_qeplus_lab_b_SA_R2.raw |
| SA | R3 | 54 | DIA_Data | 54_qeplus_lab_b_SA_R3.raw |
| SB | R1 | 54 | DIA_Data | 54_qeplus_lab_b_SB_R1.raw |
| SB | R2 | 54 | DIA_Data | 54_qeplus_lab_b_SB_R2.raw |
| SB | R3 | 54 | DIA_Data | 54_qeplus_lab_b_SB_R3.raw |
| SC | R1 | 54 | DIA_Data | 54_qeplus_lab_b_SC_R1.raw |
| SC | R2 | 54 | DIA_Data | 54_qeplus_lab_b_SC_R2.raw |
| SC | R3 | 54 | DIA_Data | 54_qeplus_lab_b_SC_R3.raw |
| SA | R1 | 54 | DIA_Data | 54_qeplus_lab_h_SA_R1.raw |
| SA | R2 | 54 | DIA_Data | 54_qeplus_lab_h_SA_R2.raw |
| SA | R3 | 54 | DIA_Data | 54_qeplus_lab_h_SA_R3.raw |
| SB | R1 | 54 | DIA_Data | 54_qeplus_lab_h_SB_R1.raw |
| SB | R2 | 54 | DIA_Data | 54_qeplus_lab_h_SB_R2.raw |
| SB | R3 | 54 | DIA_Data | 54_qeplus_lab_h_SB_R3.raw |
| SC | R1 | 54 | DIA_Data | 54_qeplus_lab_h_SC_R1.raw |
| SC | R2 | 54 | DIA_Data | 54_qeplus_lab_h_SC_R2.raw |
| SC | R3 | 54 | DIA_Data | 54_qeplus_lab_h_SC_R3.raw |
|  |  | 54 | windows.png | 54_windows-b.png |
|  |  | 54 | windows.txt | 54_windows-b.txt |
|  |  | 54 | windows.png | 54_windows-h.png |
|  |  | 54 | windows.txt | 54_windows-h.txt |
|  |  | 55 | CustomLCMethod.csv | 55_CustomLCMethod.csv |
|  |  | 55 | CustomMSMethod.csv | 55_CustomMSMethod.csv |
| SA | R1 | 55 | DIA_Data | 55_qehf_lab_SA_R1.raw |
| SB | R1 | 55 | DIA_Data | 55_qehf_lab_SB_R1.raw |
| SC | R1 | 55 | DIA_Data | 55_qehf_lab_SC_R1.raw |
|  |  | 55 | windows.png | 55_windows.png |
|  |  | 55 | windows.txt | 55_windows.txt |
|  |  | 56 | CustomMSMethod.csv | 56_CustomMSMethod.csv |
| SA | R1 | 56 | DIA_Data | 56_orbivep_lab_SA_R1.raw |
| SA | R2 | 56 | DIA_Data | 56_orbivep_lab_SA_R2.raw |
| SA | R3 | 56 | DIA_Data | 56_orbivep_lab_SA_R3.raw |
| SB | R1 | 56 | DIA_Data | 56_orbivep_lab_SB_R1.raw |
| SB | R2 | 56 | DIA_Data | 56_orbivep_lab_SB_R2.raw |
| SB | R3 | 56 | DIA_Data | 56_orbivep_lab_SB_R3.raw |
| SC | R1 | 56 | DIA_Data | 56_orbivep_lab_SC_R1.raw |
| SC | R2 | 56 | DIA_Data | 56_orbivep_lab_SC_R2.raw |
| SC | R3 | 56 | DIA_Data | 56_orbivep_lab_SC_R3.raw |
|  |  | 56 | windows.png | 56_windows.png |
|  |  | 56 | windows.txt | 56_windows.txt |
| SA | R1 | 57 | DIA_Data | 57_qehfx_prg_SA_R1.raw |
| SA | R2 | 57 | DIA_Data | 57_qehfx_prg_SA_R2.raw |
| SA | R3 | 57 | DIA_Data | 57_qehfx_prg_SA_R3.raw |
| SB | R1 | 57 | DIA_Data | 57_qehfx_prg_SB_R1.raw |
| SB | R2 | 57 | DIA_Data | 57_qehfx_prg_SB_R2.raw |
| SB | R3 | 57 | DIA_Data | 57_qehfx_prg_SB_R3.raw |
| SC | R1 | 57 | DIA_Data | 57_qehfx_prg_SC_R1.raw |
| SC | R2 | 57 | DIA_Data | 57_qehfx_prg_SC_R2.raw |
| SC | R3 | 57 | DIA_Data | 57_qehfx_prg_SC_R3.raw |
|  |  | 57 | windows.png | 57_windows.png |
|  |  | 57 | windows.txt | 57_windows.txt |
|  |  | 60 | CustomLCMethod.csv | 60_CustomLCMethod.csv |
| SA | R1 | 60 | DIA_Data | 60_lumos_lab_SA_R1.raw |
| SA | R2 | 60 | DIA_Data | 60_lumos_lab_SA_R2.raw |
| SA | R3 | 60 | DIA_Data | 60_lumos_lab_SA_R3.raw |
| SB | R1 | 60 | DIA_Data | 60_lumos_lab_SB_R1.raw |
| SB | R2 | 60 | DIA_Data | 60_lumos_lab_SB_R2.raw |
| SB | R3 | 60 | DIA_Data | 60_lumos_lab_SB_R3.raw |
| SC | R1 | 60 | DIA_Data | 60_lumos_lab_SC_R1.raw |
| SC | R2 | 60 | DIA_Data | 60_lumos_lab_SC_R2.raw |
| SC | R3 | 60 | DIA_Data | 60_lumos_lab_SC_R3.raw |
|  |  | 60 | windows.png | 60_windows.png |
|  |  | 60 | windows.txt | 60_windows.txt |
|  |  | 63 | CustomLCMethod.csv | 63_CustomLCMethod.csv |
|  |  | 63 | CustomMSMethod.csv | 63_CustomMSMethod.csv |
| SA | R1 | 63 | DIA_Data | 63_lumos_prg_SA_R1.raw |
| SA | R2 | 63 | DIA_Data | 63_lumos_prg_SA_R2.raw |
| SA | R3 | 63 | DIA_Data | 63_lumos_prg_SA_R3.raw |
| SB | R1 | 63 | DIA_Data | 63_lumos_prg_SB_R1.raw |
| SB | R2 | 63 | DIA_Data | 63_lumos_prg_SB_R2.raw |
| SB | R3 | 63 | DIA_Data | 63_lumos_prg_SB_R3.raw |
| SC | R1 | 63 | DIA_Data | 63_lumos_prg_SC_R1.raw |
| SC | R2 | 63 | DIA_Data | 63_lumos_prg_SC_R2.raw |
| SC | R3 | 63 | DIA_Data | 63_lumos_prg_SC_R3.raw |
|  |  | 63 | windows.png | 63_windows.png |
|  |  | 63 | windows.txt | 63_windows.txt |
| Pool | R1 | 03 | DDA_Data | 03_lumos_prg_DDA_pool_R1.raw |
| Pool | R2 | 03 | DDA_Data | 03_lumos_prg_DDA_pool_R2.raw |
| Pool | R3 | 03 | DDA_Data | 03_lumos_prg_DDA_pool_R3.raw |
| Pool |  | 03 | GPF_Data | 03_lumos_prg_DIA_GPF1.raw |
| Pool |  | 03 | GPF_Data | 03_lumos_prg_DIA_GPF2.raw |
| Pool |  | 03 | GPF_Data | 03_lumos_prg_DIA_GPF3.raw |
| Pool |  | 03 | GPF_Data | 03_lumos_prg_DIA_GPF4.raw |
| Pool |  | 03 | GPF_Data | 03_lumos_prg_DIA_GPF5.raw |
| Pool | R1 | 48 | DDA_Data | 48_lumos_prg_DDA_pool_R1.raw |
| Pool | R2 | 48 | DDA_Data | 48_lumos_prg_DDA_pool_R2.raw |
| Pool | R3 | 48 | DDA_Data | 48_lumos_prg_DDA_pool_R3.raw |
| Spikes |  | 48 | DDA_Data | 48_qehfx_PRGspikes_100fmol_DDA.raw |
| Spikes |  | 48 | DIA_Data | 48_qehfx_PRGspikes_100fmol_DIA.raw |
| Spikes |  | 48 | DDA_Data | 48_qehfx_PRGspikes_200fmol_DDA.raw |
| Spikes |  | 48 | DIA_Data | 48_qehfx_PRGspikes_200fmol_DIA.raw |
| Spikes |  | 48 | DDA_Data | 48_qehfx_PRGspikes_500fmol_DDA.raw |
| Spikes |  | 48 | DIA_Data | 48_qehfx_PRGspikes_500fmol_DIA.raw |
| Spikes |  | 48 | DIA_Data | 48_qehfx_PRGspikes_800fmol_DIA.raw |
| Spikes |  | 48 | DDA_Data | 48_qehfx_PRGspikess_800fmol_DDA.raw |
| Pool | R1 | 48 | DDA_Data | 48_qehfx_prg_DDA_pool_R1.raw |
| Pool | R2 | 48 | DDA_Data | 48_qehfx_prg_DDA_pool_R2.raw |
| Pool | R3 | 48 | DDA_Data | 48_qehfx_prg_DDA_pool_R3.raw |
|  |  |  | EncyclopeDIA Spectral Library | combined_prg_pan_human.dlib |
|  |  |  | EncyclopeDIA Spectral Library | combined_prg_sprothuman.dlib |
|  |  |  | FASTA | contaminants_20120713.fasta |
|  |  |  | FASTA | sp_human_180620_plus_PRG_ABRF_4_prot.fasta |
|  |  |  | Scaffold Chromatogram Library | 03 - PH plus CL library.elib |

|  |  |
| --- | --- |
| Scaffold Chromatogram Library | 03 - Prosit plus CL library.elib |
| Scaffold Chromatogram Library | 03 Chromatogram Library.elib |
| Scaffold Experiment | 03 - CL only.sdia |
| Scaffold Experiment | 03 - PH plus CL.sdia |
| Scaffold Experiment | 03 - Prosit plus CL.sdia |
| Scaffold Results Export | 03 - CL only - ABRF Spike Proteins.csv |
| Scaffold Results Export | 03 - PH plus CL - ABRF Spike Proteins.csv |
| Scaffold Results Export | 03 - Prosit plus CL - ABRF Spike Proteins.csv |
| Scaffold Results Export | Peptide Match Report of 03 - CL only.csv |
| Scaffold Results Export | Peptide Match Report of 03 - PH plus CL.csv |
| Scaffold Results Export | Peptide Match Report of 03 - Prosit plus CL.csv |
| Scaffold Results Export | Samples Report of 03 - CL only.csv |
| Scaffold Results Export | Samples Report of 03 - PH plus CL.csv |
| Scaffold Results Export | Samples Report of 03 - Prosit plus CL.csv |
| Scaffold Results Export | Samples Table of 03 - CL only.csv |
| Scaffold Results Export | Samples Table of 03 - PH plus CL.csv |
| Scaffold Results Export | Samples Table of 03 - Prosit plus CL.csv |
| Spectronaut Experiment | 03_48_against_combined_DpD.sne |
| Spectronaut Experiment | 03_48_Pan_human_Spike_lib.sne |
| Spectronaut Experiment | 03_against_48_DpD.sne |
| Spectronaut Experiment | 03_lumos_directDIA.sne |
| Spectronaut Experiment | 03_Lumos_own_DpD.sne |
| Spectronaut Experiment | 48_against_03_DpD.sne |
| Spectronaut Experiment | 48_lumos_directDIA.sne |
| Spectronaut Experiment | 48_lumos_own_DpD.sne |
| Spectronaut Export | 03_48_against_combined_DpD_new_ExperimentAnalysis.txt |
| Spectronaut Export | 03_48_against_combined_DpD_new_IdentificationsOverview.tsv |
| Spectronaut Export | 03_48_against_combined_DpD_new_PGReport.xlsx |
| Spectronaut Export | 03_48_Pan_human_Spike_lib_ExperimentAnalysis.txt |
| Spectronaut Export | 03_48_Pan_human_Spike_lib_IdentificationsOverview.tsv |
| Spectronaut Export | 03_48_Pan_human_Spike_lib_PGReport.xlsx |
| Spectronaut Export | 03_against_48_DpD_new_SA_lib_ExperimentAnalysis.txt |
| Spectronaut Export | 03_against_48_DpD_new_SA_lib_IdentificationsOverview.tsv |
| Spectronaut Export | 03_against_48_DpD_new_SA_lib_PGReport.xlsx |
| Spectronaut Export | 03_lumos_directDIA_ExperimentAnalysis.txt |
| Spectronaut Export | 03_lumos_directDIA_IdentificationsOverview.tsv |
| Spectronaut Export | 03_lumos_directDIA_PGReport.xlsx |
| Spectronaut Export | 03_Lumos_own_DpD_ExperimentAnalysis.txt |
| Spectronaut Export | 03_Lumos_own_DpD_IdentificationsOverview.tsv |
| Spectronaut Export | 03_Lumos_own_DpD_PGReport.xlsx |
| Spectronaut Export | 48_Lumos_against_DpD_03_lib_ExperimentAnalysis.txt |
| Spectronaut Export | 48_Lumos_against_DpD_03_lib_IdentificationsOverview.tsv |
| Spectronaut Export | 48_Lumos_against_DpD_03_lib_PGReport.xlsx |
| Spectronaut Export | 48_lumos_directDIA_ExperimentAnalysis.txt |
| Spectronaut Export | 48_lumos_directDIA_IdentificationsOverview.tsv |
| Spectronaut Export | 48_lumos_directDIA_PGReport.xlsx |
| Spectronaut Export | 48_lumos_own_DpD_new_ExperimentAnalysis.txt |
| Spectronaut Export | 48_lumos_own_DpD_new_IdentificationsOverview.tsv |
| Spectronaut Export | 48_lumos_own_DpD_new_PGReport.xlsx |
| Spectronaut Spectral Library | 03_lumos_48_lumos_DpD.kit |
| Spectronaut Spectral Library | 03_lumos_48_lumos_DpD.xls |
| Spectronaut Spectral Library | 03_lumos_directDIA.kit |
| Spectronaut Spectral Library | 03_lumos_directDIA.xls |
| Spectronaut Spectral Library | 03_lumos_DpD.kit |
| Spectronaut Spectral Library | 03_lumos_DpD.xls |
| Spectronaut Spectral Library | 48_lumos_directDIA.kit |
| Spectronaut Spectral Library | 48_lumos_directDIA.xls |
| Spectronaut Spectral Library | 48_lumos_DpD.kit |
| Spectronaut Spectral Library | 48_lumos_DpD.xls |
| Spectronaut Spectral Library | 48_qehfx_spikes_DpD.kit |
| Spectronaut Spectral Library | 48_qehfx_spikes_DpD.xls |
| Spectronaut Spectral Library | Pan Human Library - ETH.kit |
| Spectronaut Spectral Library | Pan Human Library - ETH.xls |
| Metadata for 49 Data Sets | Online-only Table 1.xlsx |

## Technical Validation

### Data return and curation

Participants uploaded their data to a private FTP hosted by MassIVE. Each participant was given a folder designated by their participant number and the file naming scheme was described in Supplemental File 1. The instrument naming scheme was changed for Online-only Table 1 to reflect instrument names, following the PSI-MS recommended names (https://github.com/HUPO-PSI/psi-ms-CV/blob/master/psi-ms.obo). Following the end of the study, file integrity was confirmed by opening each file. In some cases, the file was corrupt and the participant requested by the anonymiser to re-uploaded the files. In some cases, the original file was also corrupt and those data were not available. In the case of missing files, we made every effort with the participant to find and upload the missing data. Despite these efforts, not all participants were able to provide the requested nine raw data files.

Once the data was curated, files were manually inspected using the TIC to look for any noticeable qualities such as TIC without peaks. Notes were made in Online-only Table 1. The majority of replicates were consistent, though it should be noted that this does not imply a measure of data quality. Next, the embedded parameters in the raw files were used to determine and or confirm MS-acquisition settings. Though participants were encouraged to self-report MS settings, there was missing information and discrepancies. To avoid reliance on the participant, the first replicate of sample A was used to determine DIA window scheme, MS1 and MS2 resolution (and injection time if applicable) and DIA cycle time. All other files from that participant were assumed to have the same acquisition settings. In the case of Sciex TripleTOFs, participants reported MS1 and MS2 settings. The DIA windowing strategy was determined using Skyline^34^, while the scan header provided MS1 and MS2 information (in the case of Thermo instruments). The DIA cycle time was determined using Spectronaut (v13.6.190905.43655; Biognosys AG)^28^. This information is provided in Online-only Table 1.

Although LC conditions were not available from data files many participants submitted LC gradient specifics along with raw data. These are included as “supplemental” in Online-only Table 1, and are available in the MassIVE MSV000086479. Finally, in case where those metadata couldn’t be surmised and were not self-reported, we contacted the participant directly to request that information. After these efforts, there are still some participants with missing information. This is noted in Online-only Table 1.

### Survey

Participants were given the option to self-report the specific LC columns they used. the HPLC parameters, the MS instrument settings and any attempt to identify the amount of spike in proteins in the labelled samples. These survey questions are provided in Supplemental File 2.

## Usage Notes

All raw files are available on MassIVE MSV000086479, and many programs can use these files directly. Software to analyse DIA includes, but is not limited to: DIA-NN^35^, DIA-Umpire^36^, EncyclopeDIA^25^, OpenSWATH^37^, PEAKS Studio X (Bioinformatics Solutions, Inc.), PECAN^38^, Protalizer (Vulcan Analytical), Scaffold DIA (Proteome Software), Skyline^34^, and Spectronaut (Biognosys AG)^28^ as well as DIA specific statistical packages (ex. iq: Protein Quantification in Mass Spectrometry-Based Proteomics; https://CRAN.R-project.org/package=iq). Many of these programs have excellent online resources including tutorials of the analysis pipeline. The relevant information such as DIA window placement or instrument settings can be found in Online-only Table 1 and in supplemental files on MassIVE MSV000086479. Specifically to the analysis performed in this paper, we have included .sne (Spectronaut) and .sdia (Scaffold DIA) files, which are located on MassIVE MSV000086479 and can be opened with free viewers for these programs. The libraries used in the analysis can be found on MassIVE MSV000086479 as .kit, .xls, .dlib or .elib files and can be used directly by some of the software listed. Alternatively, the DDA files available on MassIVE MSV000086479 can be used to create libraries. It should be noted that all samples included the iRT peptides which can be used, if needed, to map the elution patterns into iRT space. Finally, in the case of *in silico* libraries such as Prosit^19^ and MS2PIP^21,23^, the original publication or tutorials should be consulted for instructions for how to combine with empirical data^20^.

## Code Availability

Analysis of raw data performed using Spectronaut (v13.6.190905.43655; Biognosys AG)^28^ and Scaffold DIA (v1.3.1; Proteome Software). No other code was used for this data generation or example analysis.

## Author contributions

BN assisted in study design and execution, analysed data, curated study data, wrote manuscript with contributions from all authors.

PS assisted in study design and execution, prepared experimental samples, feedback and edits on manuscript.

BCS assisted with data analysis, design and generation of figures, feedback and edits on manuscript.

LH assisted in study design and execution, design and generation of figures, feedback and edits on manuscript

LM assisted in study design and execution, feedback and edits on manuscript.

MM assisted in study design and execution, feedback and edits on manuscript.

BP assisted in study design and execution, feedback and edits on manuscript.

BS assisted in study design and execution, feedback and edits on manuscript.

MP assisted in study design and execution, feedback and edits on manuscript.

YW assisted in study design and execution, developed participant survey, feedback and edits on manuscript.

PJ assisted in study design and execution, feedback and edits on manuscript.

JK assisted in study design and execution, analysed data, wrote manuscript with contributions from all authors.

## Acknowledgements

The authors wish to thank Matt Herring for graphical advice when designing figures. Identification of certain commercial equipment, instruments, software or materials does not imply recommendation or endorsement by the National Institute of Standards and Technology, nor does it imply that the products identified are necessarily the best available for the purpose. P.D.J was supported by National Cancer Institute - Informatics Technology for Cancer Research (NCI-ITCR) grant 1U24CA199347 and National Science Foundation (U.S.) grant 1458524. We would also like to thank the software companies (Biognosys AG, Bioinformatics Solutions, Inc., and Proteome Software for providing extended trial licences of their software to study participants and Biognosys AG for the provision of the iRT peptides added to all samples.

